# Integrin alpha11 is an Osteolectin receptor and is required for the maintenance of adult skeletal bone mass

**DOI:** 10.1101/429746

**Authors:** Bo Shen, Kristy Vardy, Payton Hughes, Alpaslan Tasdogan, Zhiyu Zhao, Genevieve Crane, Sean J. Morrison

## Abstract

We previously discovered a new osteogenic growth factor that is required to maintain adult skeletal bone mass, Osteolectin/Clec11a. Osteolectin acts on Leptin Receptor^+^ (LepR^+^) skeletal stem cells and other osteogenic progenitors in bone marrow to promote their differentiation into osteoblasts. Here we identity a receptor for Osteolectin, α11 integrin, which is expressed by LepR^+^ cells and osteoblasts. α11β1 integrin binds Osteolectin with nanomolar affinity and is required for the osteogenic response to Osteolectin. Deletion of *Itga11* (which encodes α11) from mouse and human bone marrow stromal cells impaired osteogenic differentiation and blocked their response to Osteolectin. Like *Osteolectin* deficient mice, *Lepr*-*cre*; *Itga11^fl/fl^* mice appeared grossly normal but exhibited reduced osteogenesis and accelerated bone loss during aging. Osteolectin binding to α11β1 promoted Wnt pathway activation, which was necessary for the osteogenic response to Osteolectin. This reveals a new mechanism for maintenance of adult bone mass: Wnt pathway activation by Osteolectin/α11β1 signaling.

## Introduction

The maintenance of the adult skeleton requires the formation of new bone throughout life from the differentiation of skeletal stem/progenitor cells into osteoblasts. Leptin Receptor^+^ (LepR^+^) bone marrow stromal cells are the major source of osteoblasts and adipocytes in adult mouse bone marrow (Zhou et al., 2014). These cells arise postnatally in the bone marrow, where they are initially rare and make little contribution to the skeleton during development, but expand to account for 0.3% of cells in adult bone marrow (Mizoguchi et al., 2014; Zhou et al., 2014). Nearly all fibroblast colony-forming cells (CFU-F) in adult mouse bone marrow arise from these LepR^+^ cells, and a subset of LepR^+^ cells form multilineage colonies with osteoblasts, adipocytes, and chondrocytes, suggesting they are highly enriched for skeletal stem cells (Zhou et al., 2014). LepR^+^ cells are also a critical source of growth factors that maintain hematopoietic stem cells and other primitive hematopoietic progenitors in mouse bone marrow (Ding and Morrison, 2013; Ding et al., 2012; Himburg et al., 2018; Oguro et al., 2013).

To identify new growth factors that regulate hematopoiesis or osteogenesis, we performed RNA-Seq analysis on LepR^+^ cells and looked for transcripts predicted to encode secreted proteins with sizes and structures similar to growth factors and whose function had not been studied *in vivo*. We discovered that Clec11a, a secreted glycoprotein of the C-type lectin domain superfamily (Bannwarth et al., 1999; Bannwarth et al., 1998), was preferentially expressed by LepR^+^ cells (Yue et al., 2016). Prior studies had observed Clec11a expression in bone marrow but inferred based on colony-forming assays in culture that it was a hematopoietic growth factor (Hiraoka et al., 1997; Hiraoka et al., 2001). We made germline knockout mice and found it is not required for normal hematopoiesis but that it is required for the maintenance of the adult skeleton (Yue et al., 2016). The mutant mice formed their skeleton normally during development and were otherwise grossly normal as adults but exhibited reduced adult osteogenesis and accelerated bone loss during aging (Yue et al., 2016). Recombinant protein promoted osteogenic differentiation by bone marrow stromal cells *in vitro* and *in vivo* (Yue et al., 2016). Based on these observations we proposed to call this new osteogenic growth factor, Osteolectin, so as to have a name related to its biological function. Osteolectin/Clec11a is expressed by a subset of LepR^+^ stromal cells in the bone marrow as well as by osteoblasts, osteocytes, and hypertrophic chondrocytes. The discovery of Osteolectin offers the opportunity to better understand the mechanisms that maintain the adult skeleton; however, the Osteolectin receptor and the signaling mechanisms by which it promotes osteogenesis are unknown.

Several families of growth factors, and the signaling pathways they activate, promote osteogenesis, including Bone Morphogenetic Proteins (BMPs), Fibroblast Growth Factors (FGFs), Hedgehog proteins, Insulin-Like Growth Factors (IGFs), Transforming Growth Factor-betas (TGF-βs), and Wnts (reviewed by (Karsenty, 2003; Kronenberg, 2003; Wu et al., 2016). Bone marrow stromal cells regulate osteogenesis by skeletal stem/progenitor cells by secreting multiple members of these growth factor families (Chan et al., 2015). The Wnt signaling pathway is a particularly important regulator of osteogenesis, as GSK3 inhibition and β-catenin accumulation promote the differentiation of skeletal stem/progenitor cells into osteoblasts (Bennett et al., 2005; Dy et al., 2012; Hernandez et al., 2010; Krishnan et al., 2006; Kulkarni et al., 2006; Rodda and McMahon, 2006). Consistent with this, mutations that promote Wnt pathway activation increase bone mass in humans and in mice (Ai et al., 2005; Balemans et al., 2001; Boyden et al., 2002) while mutations that reduce Wnt pathway activation reduce bone mass in humans and in mice (Gong et al., 2001; Holmen et al., 2004; Kato et al., 2002).

There are 18 integrin α subunits and 8 β subunits, forming 24 different functional integrin heterodimer complexes (Humphries et al., 2006; Hynes, 1992). Integrin signaling is known to promote Wnt pathway activation through Integrin-Linked Kinase (ILK)-mediated phosphorylation of GSK3 and nuclear translocation of β-catenin (Burkhalter et al., 2011; Delcommenne et al., 1998; Novak et al., 1998; Rallis et al., 2010). Conditional deletion of *ILK* or *Focal Adhesion Kinase* (*FAK*) from osteoblast progenitors reduces osteogenesis and depletes trabecular bone in adult mice (Dejaeger et al., 2017; Sun et al., 2016), suggesting a role for integrins in adult osteogenesis. Conditional deletion of β1 integrin from chondrocytes or skeletal stem/progenitor cells impairs chondrocyte function and skeletal ossification during development (Aszodi et al., 2003; Raducanu et al., 2009; Shekaran et al., 2014). Activation of αvβ1 signaling by Osteopontin (Chen et al., 2014) or α5β1 signaling by Fibronectin (Hamidouche et al., 2009; Moursi et al., 1997) promotes the osteogenic differentiation of mesenchymal progenitors. Germline deletion of integrin α10 leads to defects in chondrocyte proliferation and growth plate function (Bengtsson et al., 2005) and germline deletion of integrin α11 leads to defects in tooth development (Popova et al., 2007). However, little is known about which integrins are required for adult osteogenesis *in vivo*.

## Results

### Integrin α11 is selectively expressed by LepR+ cells and osteoblasts

The *Osteolectin/Clec11a* gene first appeared in bony fish and is conserved among bony vertebrates (Yue et al., 2016). Osteolectin contains a glutamic acid-rich sequence, an alpha-helical leucine zipper, and a C-type lectin domain (Figure 1A and 1B) (Bannwarth et al., 1999; Bannwarth et al., 1998). To generate hypotheses regarding potential Osteolectin receptors, we examined the amino acid sequence and found two integrin-binding motifs, RGD (Gardner and Hynes, 1985; Pierschbacher and Ruoslahti, 1984; Plow et al., 1985) and LDT (Fong et al., 1997; Viney et al., 1996) in human (Figure 1A) and mouse Osteolectin (Figure 1B). One or both of these motifs were conserved across Osteolectin sequences in all bony vertebrates (Figure 1C). This raised the possibility that the Osteolectin receptor might be an integrin.

**Figure 1.**
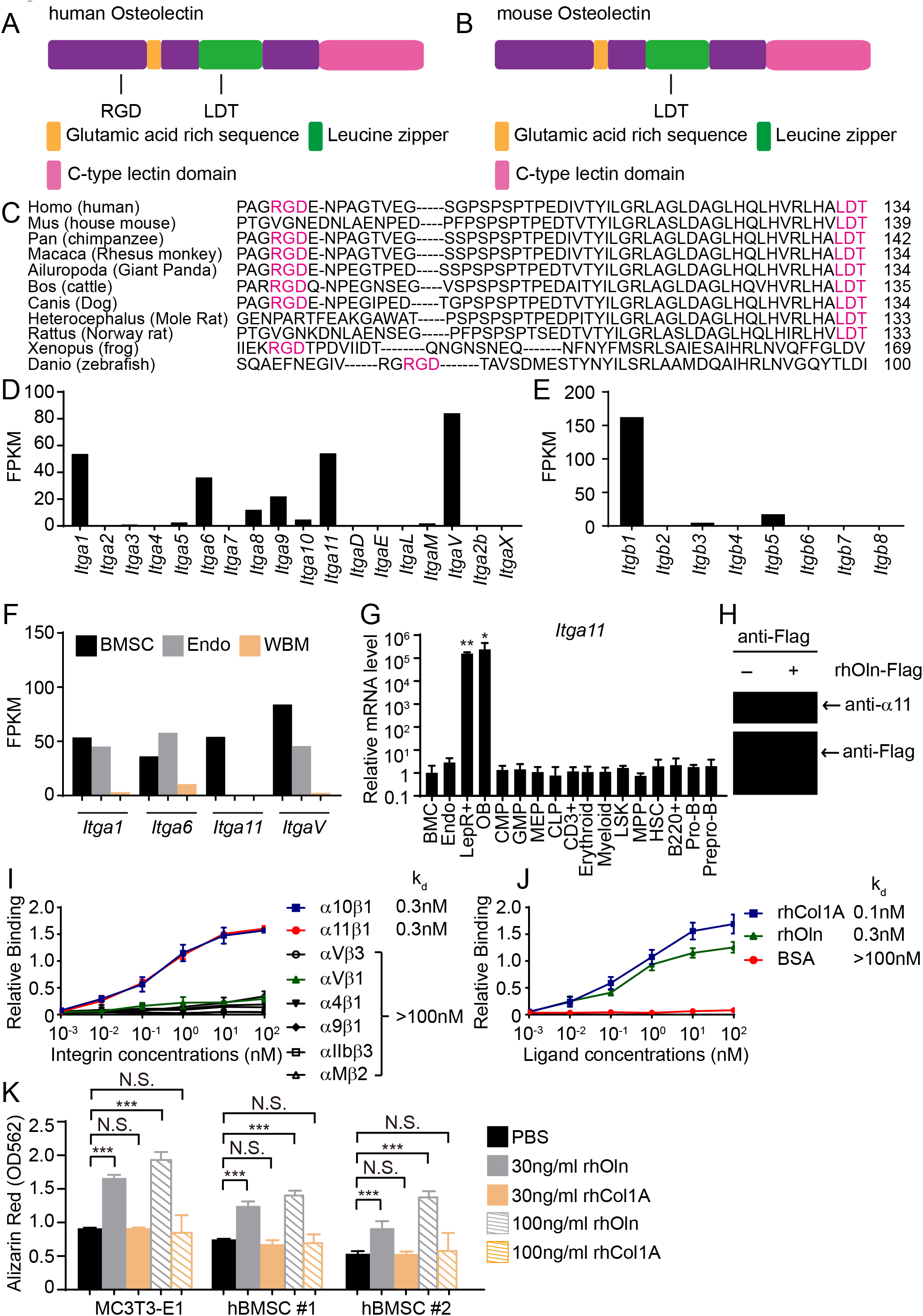
Osteolectin contains conserved integrin binding motifs and binds with high affinity to integrin α_11_β_1_. **(A, B)** The human (**A**) and mouse (**B**) Osteolectin proteins contain RGD and LDT sequences. **(C)** Alignment of Osteolectin amino acid sequences shows that the RGD and LDT domains are evolutionarily conserved among bony vertebrates. **(D, E)** RNA-seq analysis of integrin α (**D**) and β (**E**) subunits in PDGFRα^+^CD45^-^Ter119^-^CD31^-^ bone marrow stromal cells from enzymatically dissociated adult bone marrow (n = 2 independent samples). These cells are uniformly positive for LepR expression (Zhou et al., 2014). (**F**) RNA-seq analysis of *Itga1, Itga6, Itga11*, and *ItgaV* in PDGFRα^+^CD45^-^Ter119^-^CD31^-^ bone marrow stromal cells, VE-Cadherin^+^ bone marrow endothelial cells, and whole bone marrow cells (n = 2 independent samples per cell population). (**G**) *Itga11* expression in cell populations from mouse bone marrow by qRT-PCR (n = 3 independent samples per cell population). The markers used for the isolation of each cell population are shown in Supplemental Table 1. (**H**) In MC3T3-E1 preosteoblast cells expressing Flag-tagged Osteolectin, anti-Flag antibody co-immunoprecipitated endogenous integrin α_11_ with Flag-tagged Osteolectin (results are representative of two independent experiments). (**I**) Recombinant human Osteolectin (rhOln) selectively bound to recombinant human integrin α_11_β_1_ and α_10_β_1_, but not to other integrins (n = 3 independent experiments). (**J**) Integrin α_11_β_1_ bound Osteolectin and recombinant human Pro-Collagen 1α (rhCol1A) with similar affinities, but not bovine serum albumin (BSA) (n = 3 independent experiments). (**K**) Osteolectin, but not Pro-Collagen 1α, promoted osteogenic differentiation by MC3T3-E1 cells and human bone marrow stromal cells (n = 3 independent experiments). All numerical data reflect mean ± standard deviation. Statistical significance was determined with one-way (**G**) or two-way (**K**) ANOVAs with Dunnett’s multiple comparisons tests.

We hypothesized that the Osteolectin receptor would be restricted in its expression to osteogenic cells in the bone marrow, so we examined the expression of all α and β integrins in mouse bone marrow stromal cells by RNA-seq analysis. Among the genes that encode α integrins, *Itga1* (encoding α1), *Itga6* (encoding α6), *Itga11* (encoding α11) and *ItgaV* (encoding αV), were strongly expressed by bone marrow stromal cells (Figure 1D). Among the genes that encode β integrins, only *Itgb1* (encoding β1) was strongly expressed by bone marrow stromal cells (Figure 1E). To refine potential candidate receptors for Osteolectin we more closely examined the expression of these genes in LepR^+^ cells, which are osteogenic and should express the receptor, and endothelial cells, which are not osteogenic and should not express the receptor. *Itga1, Itga6*, and *ItgaV* were strongly expressed by both LepR^+^ cells and endothelial cells, suggesting they did not encode the receptor (Figure 1F). *Itga11* was expressed exclusively by LepR^+^ cells, not by endothelial cells or other bone marrow cells (Figure 1F). We next examined *Itga11* expression in a number of sorted bone marrow cell populations by quantitative reverse transcription PCR (qRT-PCR) (Figure S1 shows the markers used to isolate these cell populations). We found that *Itga11* was highly expressed by LepR^+^CD45^-^Ter119^-^CD31^-^ stromal cells and *Col2.3*-*GFP*^+^CD45^-^Ter119^-^CD31^-^ osteoblasts but not by any hematopoietic stem or progenitor population (Figure 1G). The expression patterns of *Itga11* and *Itgb1* are thus consistent with a potential role in osteogenesis.

Consistent with our results, integrin α11 is expressed by human bone marrow stromal cells in a way that correlates with osteogenic potential in culture (Kaltz et al., 2010); however, α11 is not known to regulate osteogenesis. Integrin α11 heterodimerizes with integrin β1 (Velling et al., 1999) and the only known ligand for α11β1 is collagen (Popova et al., 2004; Velling et al., 1999). Few cells express integrin α11 and it has been studied less than other integrins. *Itga11* deficient mice are growth retarded and have smaller bones, but this was thought to be a consequence of a defect in incisor development that leads to malnutrition (Popova et al., 2007).

### Osteolectin binds to α11β1 with nanomolar affinity

To begin to test whether α11β1 binds Osteolectin, we overexpressed human Osteolectin with a C-terminal Flag tag in MC3T3-E1 pre-osteoblast cells and immunoprecipitated with anti-Flag beads. The anti-Flag beads pulled down the tagged Osteolectin along with endogenous integrin α11 (Figure 1H). We then tested the affinity of recombinant human Osteolectin for multiple recombinant human integrin complexes by a microtiter well binding assay. Osteolectin selectively bound to integrin α11β1 and α10β1, but not to other integrins, including αVβ1, αVβ3, α4β1, α9β1, αIIbβ3 or αMβ3 (Figure 1I). Integrin α10 is the gene most closely related to α11. Integrin α10 is expressed by osteoblasts and chondrocytes (Bengtsson et al., 2005; Engel et al., 2013) but only at a low level by bone marrow stromal cells (Figure 1D). The dissociation constant (k_d_) of Osteolectin for α10β1 and α11β1 was 0.3nM whereas the k_d_ of Osteolectin for other integrins was > 100nM. The k_d_ of human Pro-Collagen 1α for α11β1 was also high (0.1nM; Figure 1J); however, in contrast to Osteolectin, addition of Pro-Collagen 1a to culture did not promote osteogenesis by MC3T3-E1 cells or two primary human bone marrow stromal cell lines (hBMSC#1 or hBMSC#2; Figure 1K).

### Osteolectin promotes Wnt pathway activation

MC3T3-E1 cells, hBMSC#1 cells, and hBMSC#2 cells secrete Osteolectin into the culture medium (Figure 2A), consistent with our observation that Osteolectin is synthesized by a subset of LepR^+^ bone marrow stromal cells (Yue et al., 2016). Deletion of *Osteolectin* from these cell lines reduced their osteogenic differentiation in osteogenic differentiation medium (Figure 2B), demonstrating that autocrine Osteolectin production is part of what drives osteogenesis by these cells in culture.

**Figure 2.**
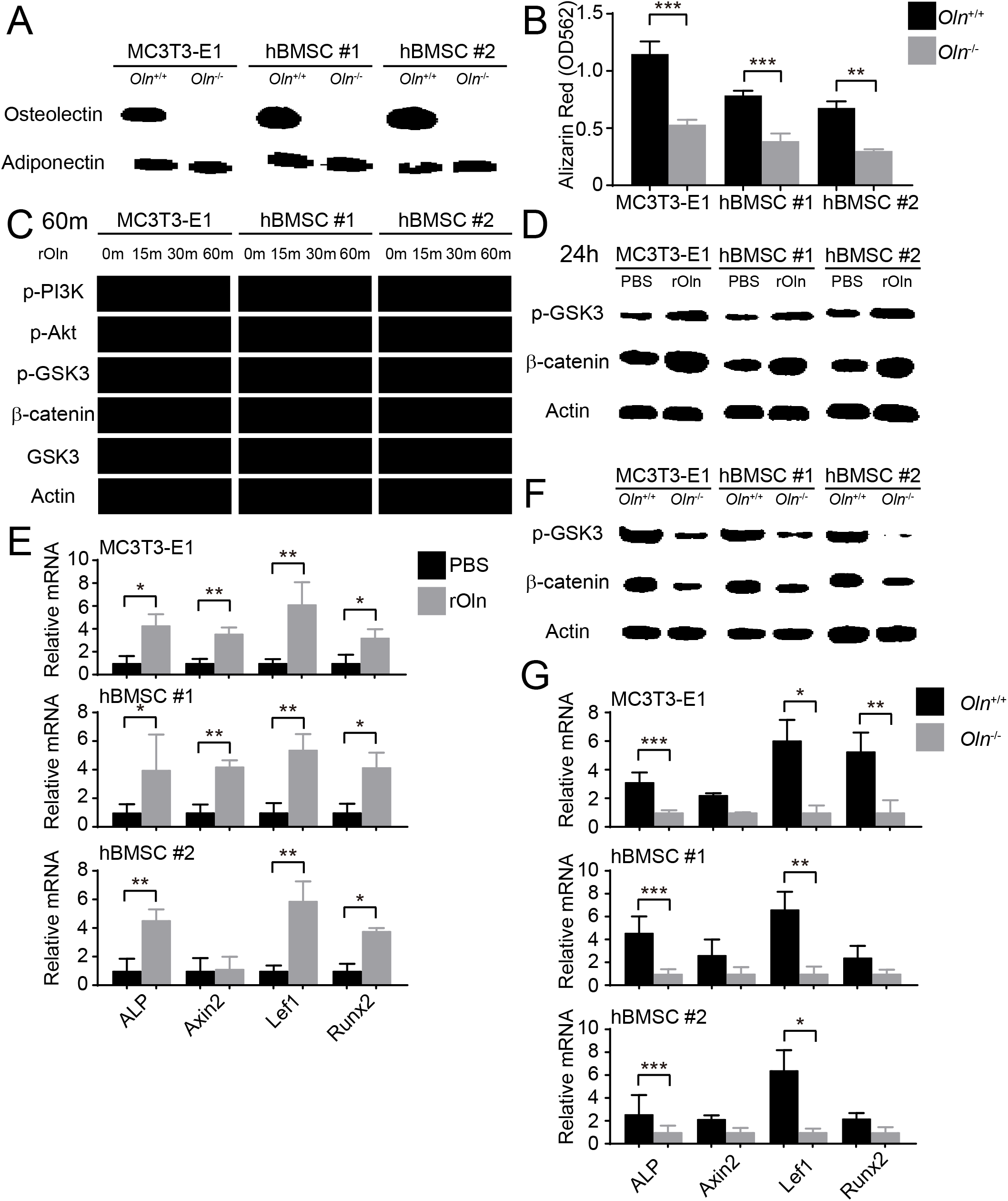
Osteolectin activates Wnt pathway signaling in skeletal stem/progenitor cells. **(A)** Western blot of cell culture supernatant from parental or *Osteolectin* deficient MC3T3-E1 cells and human bone marrow stromal cells (hBMSC#1 and hBMSC#2 cells). **(B)** Osteogenic differentiation in culture of parental or *Osteolectin* deficient MC3T3-E1 cells, hBMSC#1 cells, and hBMSC#2 cells. Alizarin red staining was performed after 14 days (for MC3T3-E1 cells) or 21 days (for hBMSC cells) to quantify osteoblast differentiation and mineralization (n = 3 independent experiments). **(C)** MC3T3-E1 cells, hBMSC#1 cells, and hBMSC#2 cells were stimulated with recombinant mouse Osteolectin (rOln) in osteogenic differentiation medium then lysed 0, 15, 30, or 60 minutes later and immunoblotted for phospho-PI3K, phospho-Akt, phospho-GSK3, β-catenin, total GSK3, and Actin (results are representative of 3 independent experiments). **(D)** MC3T3-E1 cells, hBMSC#1 cells, and hBMSC#2 cells were transferred into osteogenic differentiation medium for 24 hours with PBS or recombinant mouse Osteolectin, lysed, and immunoblotted for phospho-GSK3, β-catenin, and Actin. **(E)** qPCR analysis of Wnt target gene transcript levels (*Alkaline phosphatase, Axin2, Lef1*, or *Runx2*) in MC3T3-E1 cells, hBMSC#1 cells, and hBMSC#2 cells 24 hours after transfer into osteogenic differentiation medium, with PBS or recombinant mouse Osteolectin (n = 3 independent experiments). **(F)** Parental or *Osteolectin* deficient MC3T3-E1 cells, hBMSC#1 cells, and hBMSC#2 cells after 24 hours in osteogenic differentiation medium were lysed and immunoblotted for phospho-GSK3, β-catenin, and Actin. **(G)** qPCR analysis of Wnt target gene transcript levels in parental or *Osteolectin* deficient MC3T3-E1 cells, hBMSC#1 cells, and hBMSC#2 cells 24 hours after transfer into osteogenic differentiation medium (n = 3 independent experiments). All numerical data reflect mean ± standard deviation. The statistical significance of differences was determined with two-way ANOVAs with Sidak’s multiple comparisons tests.

To assess the signaling mechanisms by which Osteolectin promotes osteogenesis, we treated parental MC3T3-E1 cells, hBMSC#1 cells, and hBMSC#2 cells with recombinant Osteolectin and assessed the levels of phosphorylated PI3-kinase, Akt, and GSK3. The most prominent change we observed was a dramatic increase of GSK3 phosphorylation at Ser21/9 within 30 to 60 minutes of Osteolectin treatment in all three cell lines (Figure 2C). Phosphorylation at Ser21/9 inhibits the GSK3-mediated degradation of β-catenin, increasing β-catenin levels and promoting the transcription of Wnt pathway target genes (Cross et al., 1995; Peifer et al., 1994; Yost et al., 1996). We did not observe an increase in β-catenin levels within 1 hour of Osteolectin treatment, but did detect increased b-catenin levels in all three cell lines within 24 hours of Osteolectin treatment (Figure 2D). The transcription of several Wnt target genes, including *Axin2* (Jho et al., 2002; Lustig et al., 2002; Yan et al., 2001; Yan et al., 2009), *Lef1* (Filali et al., 2002; Gaur et al., 2005; Hovanes et al., 2001), *Runx2* (Dong et al., 2006; Gaur et al., 2005), and *Alkaline phosphatase* (Rawadi et al., 2003), were activated within 24 hours of Osteolectin treatment (Figure 2E). *Osteolectin* deficient MC3T3-E1 cells, hBMSC#1 cells, and hBMSC#2 cells had lower levels of phospho-GSK3 and β-catenin as compared to parental cells (Figure 2F) as well as significantly lower levels of Wnt target genes (Figure 2G). These data demonstrate that Osteolectin promotes Wnt pathway activation in osteogenic cells.

### Osteogenesis in response to Osteolectin requires β-catenin

To test whether Wnt pathway activation phenocopies the effects of Osteolectin, we evaluated the effects of AZD2858, which inhibits GSK3 function and promotes β-catenin accumulation (Berg et al., 2012). As expected (Gilmour et al., 2013; Marsell et al., 2012; Sisask et al., 2013), AZD2858 increased GSK3 phosphorylation and β-catenin levels (Figure 3A) as well as osteogenic differentiation (Figure 3B) by MC3T3-E1 cells, hBMSC#1 cells, and hBMSC#2 cells. Osteolectin also increased GSK3 phosphorylation, β-catenin levels, and osteogenic differentiation by MC3T3-E1 cells, hBMSC#1 cells, and hBMSC#2 cells (Figure 3A and 3B). However, when the two agents were added together, there was no further promotion of osteogenic differentiation beyond the effects of the individual agents (Figure 3B). These data suggest that Osteolectin and GSK3/β-catenin act in the same pathway to promote osteogenic differentiation by mesenchymal progenitors.

**Figure 3.**
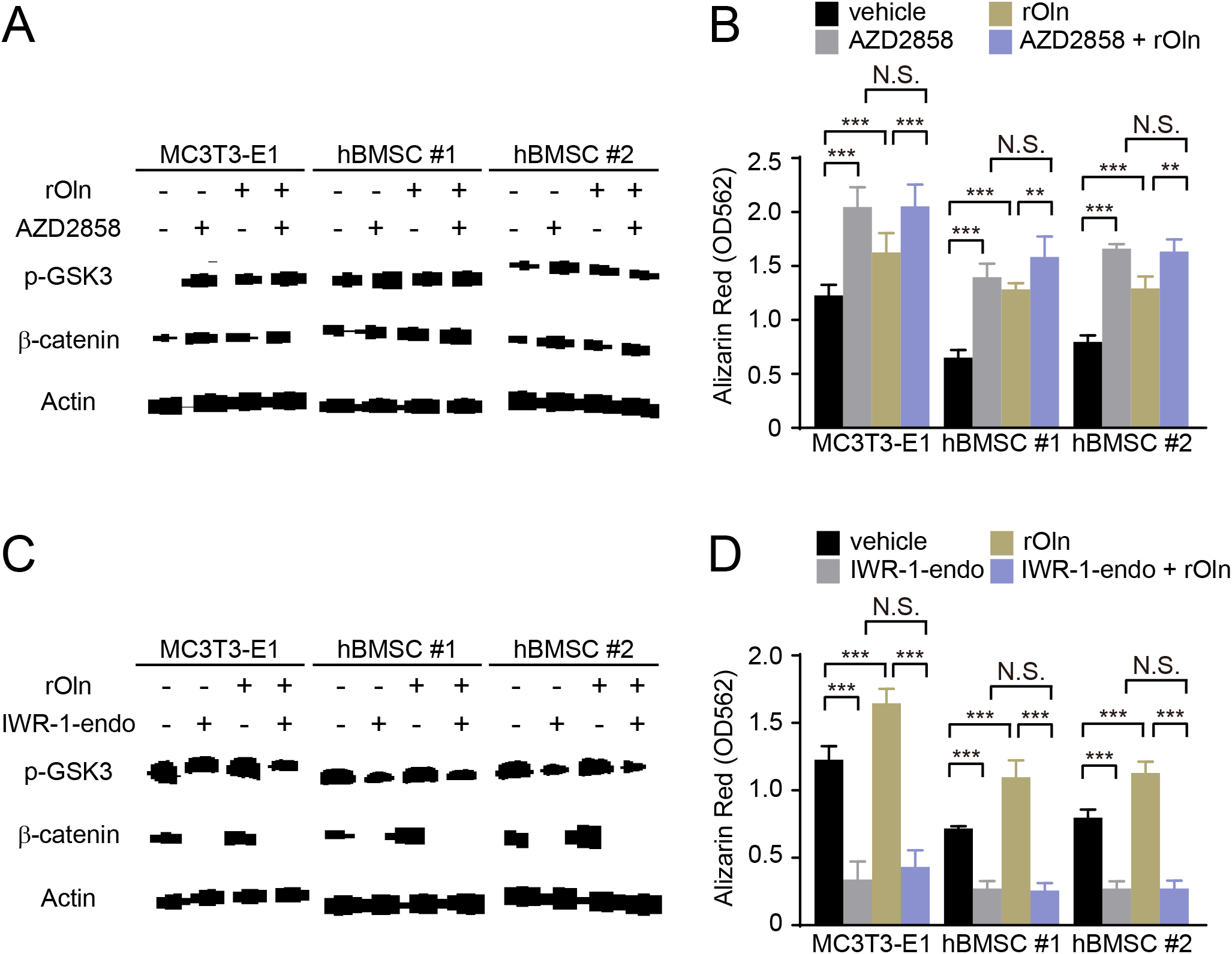
Osteogenic differentiation in response to Osteolectin requires β-catenin. (**A** and **B**) MC3T3-E1 cells, hBMSC#1 cells, and hBMSC#2 cells were transferred into osteogenic differentiation medium with PBS or 30 ng/ml recombinant mouse Osteolectin, as well as DMSO or 200 nM of the GSK3 inhibitor AZD2858. (**A**) Cells were lysed 24 hours later and immunoblotted for phospho-GSK3, β-catenin, and Actin. (**B**) Alizarin red staining after 14 days (MC3T3-E1 cells) or 21 days (hBMSC cells) to quantify osteoblast differentiation and mineralization (n = 3 independent experiments). (**C** and **D**) MC3T3-E1 cells, hBMSC#1 cells, and hBMSC#2 cells were transferred into osteogenic differentiation medium with PBS or 30 ng/ml Osteolectin, as well as DMSO or 200 nM of the β-catenin inhibitor IWR-1-endo. (**C**) Cells were lysed 24 hours later and immunoblotted for phospho-GSK3, β-catenin, and Actin. (**D**) Alizarin red staining after 14 days (MC3T3-E1 cells) or 21 days (hBMSC cells) to quantify osteoblast differentiation and mineralization (n = 3 independent experiments). All numerical data reflect mean ± standard deviation. The statistical significance of differences was determined with twoway ANOVAs with Tukey’s multiple comparisons tests.

To test whether Osteolectin requires β-catenin to promote osteogenesis, we evaluated an inhibitor of Wnt pathway signaling, IWR-1-endo, which depletes β-catenin by stabilizing Axin2 in the β-catenin destruction complex (Chen et al., 2009). As expected, Osteolectin increased β-catenin levels and osteogenesis by MC3T3-E1 cells, hBMSC#1 cells, and hBMSC#2 cells while IWR-1-endo reduced β-catenin levels and osteogenesis (Figure 3C and 3D). When added together, IWR-1-endo blocked the effect of Osteolectin on β-catenin levels and osteogenesis (Figure 3C and 3D). This suggests that Osteolectin requires β-catenin to induce osteogenesis by mesenchymal progenitors.

### Wnt pathway activation and osteogenesis by Osteolectin require α11 integrin

To test if Integrin α11 is required for the osteogenic response to Osteolectin, we used CRISPR/Cas9 to delete *Itga11* from MC3T3-E1 cells, hBMSC#1 cells, and hBMSC#2 cells. Deletion of *Itga11* from MC3T3-E1 cells, hBMSC#1 and hBMSC#2 cells significantly reduced osteogenesis by each cell line in culture (Figure 4A). Addition of recombinant Osteolectin to culture significantly promoted osteogenesis by parental, but not *Itga11* deficient, MC3T3-E1, hBMSC#1, and hBMSC#2 cells (Figure 4A). Integrin α11 is therefore required by mouse and human bone marrow stromal cells to undergo osteogenesis in response to Osteolectin.

**Figure 4.**
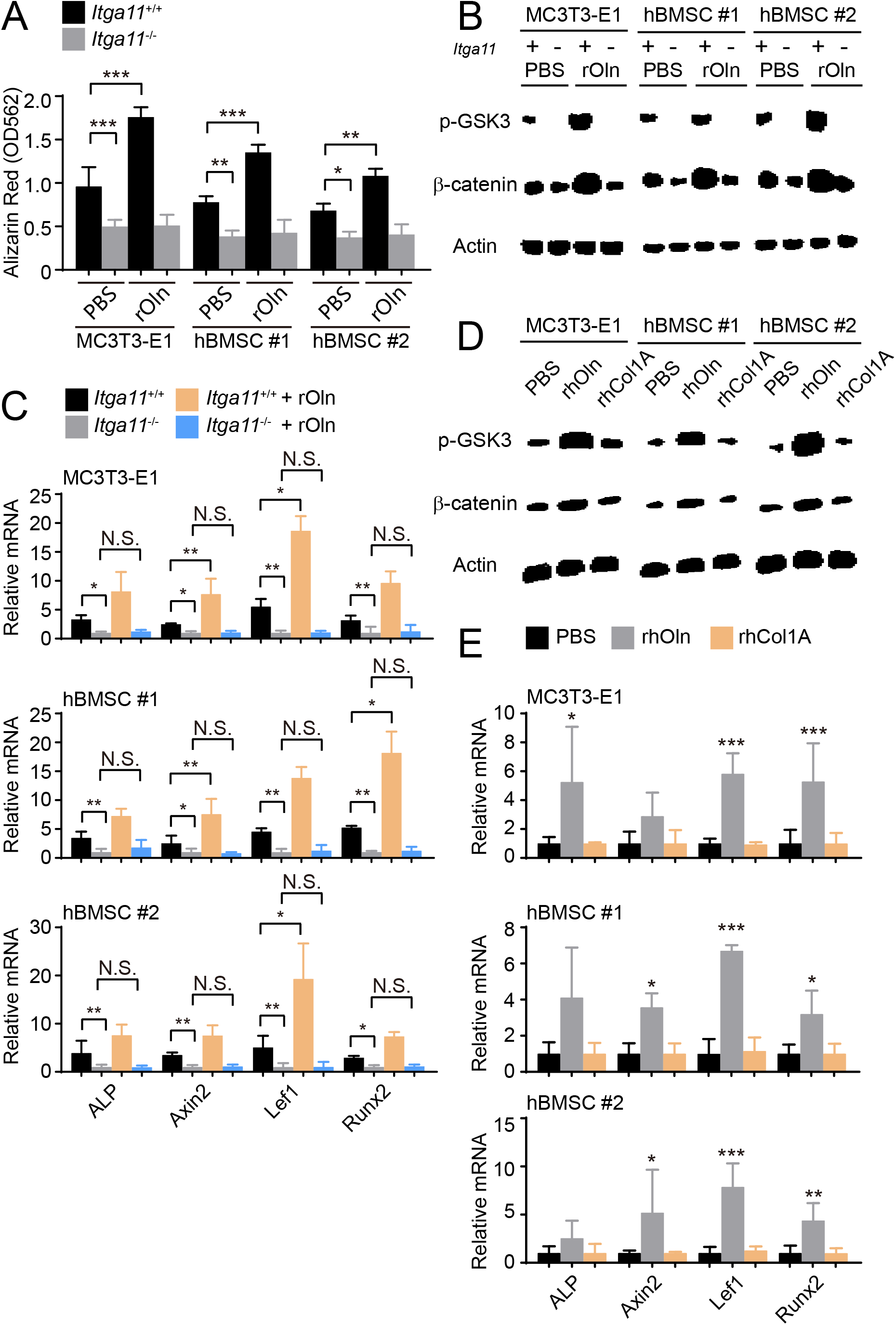
Integrin α11 is required for osteogenic differentiation and Wnt pathway activation in response to Osteolectin but exogenous collagen does not activate Wnt pathway signaling. (**A**) Osteogenic differentiation in culture of parental or *Itga11* deficient MC3T3-E1 cells, hBMSC#1 cells, and hBMSC#2 cells with PBS or recombinant mouse Osteolectin (n = 3 independent experiments). (**B**) Parental or *Itga11* deficient MC3T3-E1 cells, hBMSC#1 cells, and hBMSC#2 cells were transferred into osteogenic differentiation medium with or without recombinant mouse Osteolectin then lysed 24 hours later and immunoblotted for phospho-GSK3, β-catenin, and Actin. (**C**) qPCR analysis of Wnt target gene transcript levels in parental or *Itga11* deficient MC3T3-E1 cells, hBMSC#1 cells, and hBMSC#2 cells 24 hours after transfer into osteogenic differentiation medium with PBS or Osteolectin (n = 3 independent experiments). (**D**) MC3T3-E1 cells, hBMSC#1 cells, and hBMSC#2 cells were transferred into osteogenic differentiation medium with PBS or 30 ng/ml Osteolectin, or 30 ng/ml recombinant Pro-Collagen 1α, then lysed 24 hours later and immunoblotted for phospho-GSK3, β-catenin, and Actin. (**E**) qPCR analysis of Wnt target gene transcript levels in MC3T3-E1 cells, hBMSC#1 cells, and hBMSC#2 cells 24 hours after transfer into osteogenic differentiation medium with PBS or 30 ng/ml Osteolectin, or 30 ng/ml Pro-Collagen 1α (n = 3 independent experiments). All numerical data reflect mean ± standard deviation. Statistical significance was determined with two-way ANOVAs with Tukey’s multiple comparisons tests (**A** and **C**) or Dunnett’s multiple comparisons tests (**E**).

The *Itga11* deficient MC3T3-E1 cells, hBMSC#1 cells, and hBMSC#2 cells also exhibited lower levels of GSK3 phosphorylation and β-catenin as compared to parental control cells (Figure 4B). Addition of recombinant Osteolectin to culture increased the levels of phosphorylated GSK3 and β-catenin in parental control cells but not in *Itga11* deficient cells (Figure 4B). Consistent with the reduced levels of phosphorylated GSK3 and β-catenin in the *Itga11* deficient cells, we also observed significantly lower levels of Wnt target gene transcripts in the *Itga11* deficient cells (Figure 4C). Addition of recombinant Osteolectin to culture significantly increased the levels of Wnt target gene transcripts in parental control cells but not in *Itga11* deficient cells (Figure 4C). Integrin α11 is therefore required by mouse and human bone marrow stromal cells to activate Wnt pathway signaling in response to Osteolectin.

Given that Collagen 1 α bound to α11β1 (Figure 1J) but did not promote osteogenesis by MC3T3-E1 cells, hBMSC#1 cells, or hBMSC#2 cells (Figure 1K), we tested whether Collagen 1α promoted Wnt pathway activation in these cells. Addition of recombinant Osteolectin to culture increased levels of phosphorylated GSK3, β-catenin, and Wnt target gene transcripts in MC3T3-E1 cells, hBMSC#1 cells, and hBMSC#2 cells (Figure 4D and 4E); however, addition of Pro-Collagen 1 α to these cells did not seem to have any effect on the levels of phosphorylated GSK3, β-catenin, or Wnt target gene transcripts (Figure 4D and 4E). Exogenous Pro-Collagen 1α, therefore, did not activate Wnt pathway signaling in mouse or human bone marrow stromal cells, offering a potential explanation for its failure to promote osteogenesis by these cells.

### Conditional deletion of Integrin α11 from LepR^+^ cells reduces osteogenesis *in vivo*

To test whether Integrin α11 is necessary for osteogenesis *in vivo* we generated mice bearing a floxed allele of *Itga11* (Figure S5A-C), then conditionally deleted it from skeletal stem and progenitor cells in the bone marrow using *Lepr*-Cre. *Lepr*-*Cre*; *Itga11*^fl/fl^ mice did not exhibit the defects in incisor development (data not shown) or the growth retardation observed in germline *Itga11*^-/-^ mice (Popova et al., 2007). *Lepr*-*Cre*; *Itga11*^fl/fl^ mice appeared grossly normal (Figure 5A), with body lengths (Figure 5B), body masses (Figure 5C), and femur lengths (Figure 5D) that did not significantly differ from sex-matched littermate controls. Serum Osteolectin levels did not significantly differ between *Lepr*-*Cre*; *Itga11*^fl/fl^ mice and littermate controls at 2 or 12 months of age but were modestly higher in *Lepr*-*Cre*; *Itga11*^fl/fl^ mice as compared to controls at 6 months of age (Figure 5E). This demonstrates that Integrin α11 is not required for the synthesis or secretion of Osteolectin.

**Figure 5.**
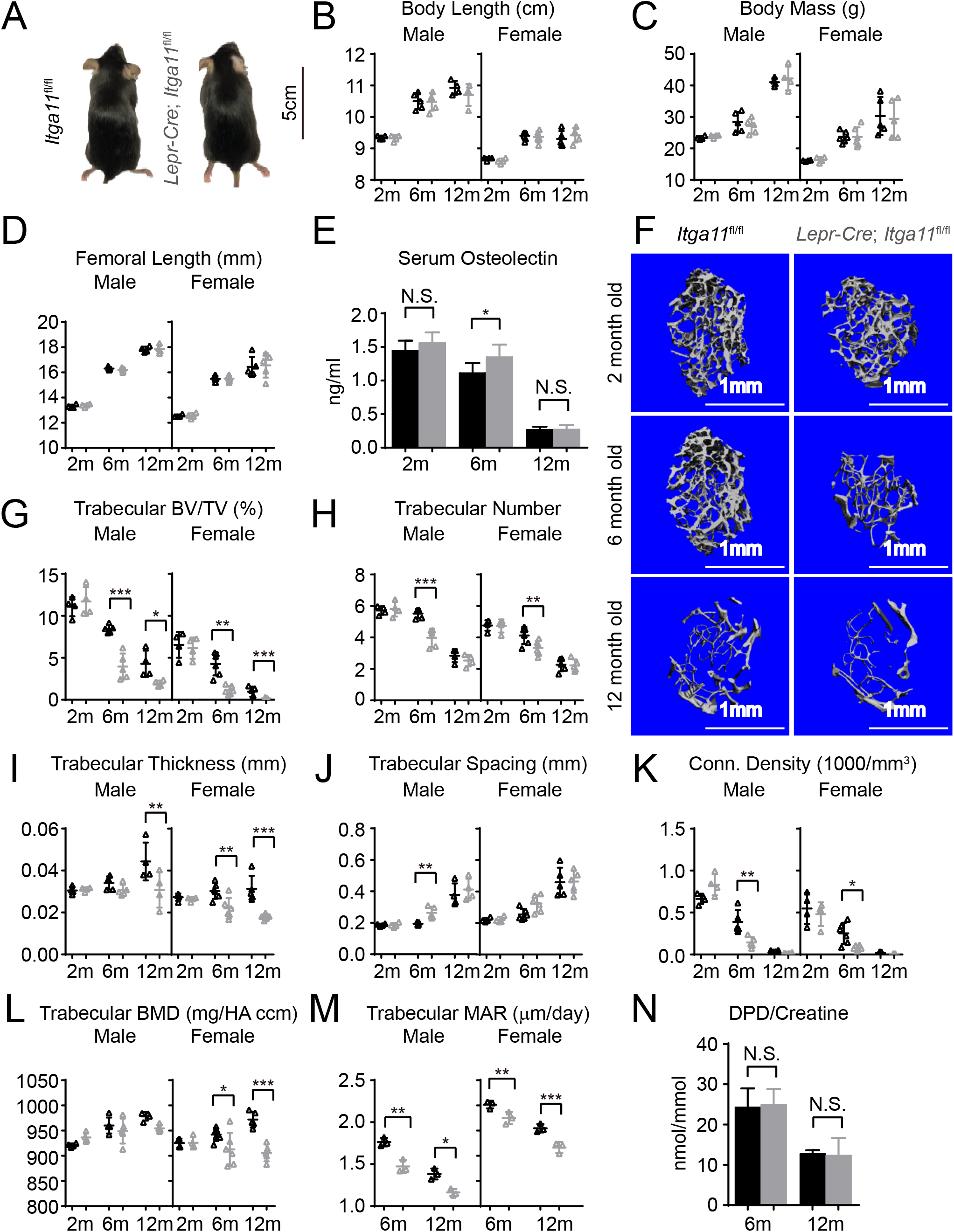
Conditional *Itga11* deletion from LepR^+^ cells accelerates the loss of trabecular bone during aging. (**A**) *Lepr*-*Cre*; *Itga11^fl/fl^* mice were grossly normal and indistinguishable from littermate controls. (**B** – **D**) Body length (**B**), body mass (**C**) and femur length (**D**) did not significantly differ between *Lepr*-*Cre*; *Itga11^fl/fl^* mice and sex-matched littermate controls at 2, 6, or 12 months of age (n = 4–6 mice per genotype per sex per time point, from at least three independent experiments). (**E**) ELISA measurement of serum Osteolectin levels in *Lepr*-*Cre*; *Itga11^fl/fl^* mice and littermate controls at 2, 6, and 12 months of age (n = 8 mice per genotype per time point from four independent experiments). (**F**) Representative microCT images of trabecular bone in the distal femur metaphysis of male *Lepr*-*Cre*; *Itga11^fl/fl^* mice and littermate controls at 2, 6, and 12 months of age. (**G**–**L**) microCT analysis of trabecular bone volume/total volume (**G**), trabecular number (**H**), trabecular bone thickness (**I**), trabecular bone spacing (**J**), connectivity density (**K**), and bone mineral density (**L**) in the distal femur metaphysis of *Lepr-Cre*; *Itga11^fl/fl^* mice and sex-matched littermate controls at 2, 6, and 12 months of age (n = 4–6 mice per genotype per sex per time point from at least three independent experiments). (**M**) Trabecular bone mineral apposition rate based on calcein double labelling in the distal femur metaphysis (n = 3 mice per genotype per sex per time point). (**N**) Bone resorption rate analysis by measuring the deoxypyridinoline/creatinine ratio in the urine of *LepR*-*Cre*; *Itga11^fl/fl^* mice and littermate controls at 6 and 12 months of age (n = 4–5 mice per genotype per time point from three independent experiments). All numerical data reflect mean ± standard deviation. The statistical significance of differences was determined with two-way ANOVAs with Sidak’s multiple comparisons tests.

To test whether deletion of *Itga11* from LepR^+^ cells affected osteogenesis in vivo, we performed micro-CT analysis of the distal femur from 2, 6, and 12-month old *Lepr*-*Cre*; *Itga11*^fl/fl^ mice and sex-matched littermates. Consistent with the observation that LepR^+^ cells and Osteolectin contribute little to skeletal development (Yue et al., 2016; Zhou et al., 2014), we observed no significant difference in trabecular bone parameters between *Lepr*-*Cre*; *Itga11*^fl/fl^ mice and sex-matched littermates at 2 months of age (Figure 5F-L). However, LepR^+^ cells and Osteolectin are necessary for adult osteogenesis (Yue et al., 2016; Zhou et al., 2014). Consistent with this, 6 and 12-month-old male and female *Lepr*-*Cre*; *Itga11*^fl/fl^ mice had significantly reduced trabecular bone volume as compared to sex-matched littermate controls (Figure 5F and 5G). At 6 and 12 months of age, male and female *Lepr*-*Cre*; *Itga11*^fl/fl^ mice also tended to have significantly lower trabecular number (Figure 5H) and trabecular thickness (Figure 5I) than sex matched littermate controls. Calcein double labelling showed that the mineral apposition rate was significantly reduced in trabecular bone from *Lepr*-*Cre*; *Itga11*^fl/fl^ mice as compared to sex-matched littermates at 6 and 12 months of age (Figure 5M). Integrin α11 is, therefore, required by LepR^+^ cells and their progeny for normal rates of trabecular bone formation and maintenance of trabecular bone mass during adulthood, phenocopying the accelerated trabecular bone loss in adult *Osteolectin* deficient mice (Yue et al., 2016).

While *Lepr*-*Cre*; *Itga11*^fl/fl^ mice had significantly reduced rates of bone formation as compared to sex-matched littermates (Figure 5M), they did not significantly differ in the urinary bone resorption marker deoxypyridinoline at 6 or 12 months of age (Figure 5N; this was not tested in 2-month-old mice because no difference in bone parameters was observed at that age). This suggests that, like Osteolectin, Integrin α11 promotes bone formation but does not regulate bone resorption.

*Osteolectin* deficiency has a milder effect on cortical bone as compared to trabecular bone, with no significant reduction in cortical bone until after 10 months of age (Yue et al., 2016). Consistent with this, femur cortical bone parameters did not significantly differ between *Lepr*-*Cre*; *Itga11*^fl/fl^ mice and sex-matched littermates at 2 or 6 month of age (Figure 6A-F). However, cortical thickness, cortical area, and cortical bone mineral density were significantly lower in male *Lepr*-*Cre*; *Itga11*^fl/fl^ mice as compared to sex-matched littermates at 12 months of age (Figure 6D-F). The differences in these parameters between female *Lepr*-*Cre*; *Itga11*^fl/fl^ mice and littermate controls were not statistically significant (Figure 6D-F). Calcein double labelling revealed that the mineral apposition rate was significantly reduced in cortical bone from male and female *Lepr*-*Cre*; *Itga11*^fl/fl^ mice as compared to sex-matched littermate controls at 6 and 12 months of age (Figure 6G). Deletion of Integrin α11 from LepR^+^ cells thus reduces the rate of cortical bone formation during adulthood, slowly leading to a thinning of cortical bone that became apparent in the femurs of male mice at 12 months of age, phenocopying the slow loss of cortical bone in adult *Osteolectin* deficient mice (Yue et al., 2016).

**Figure 6.**
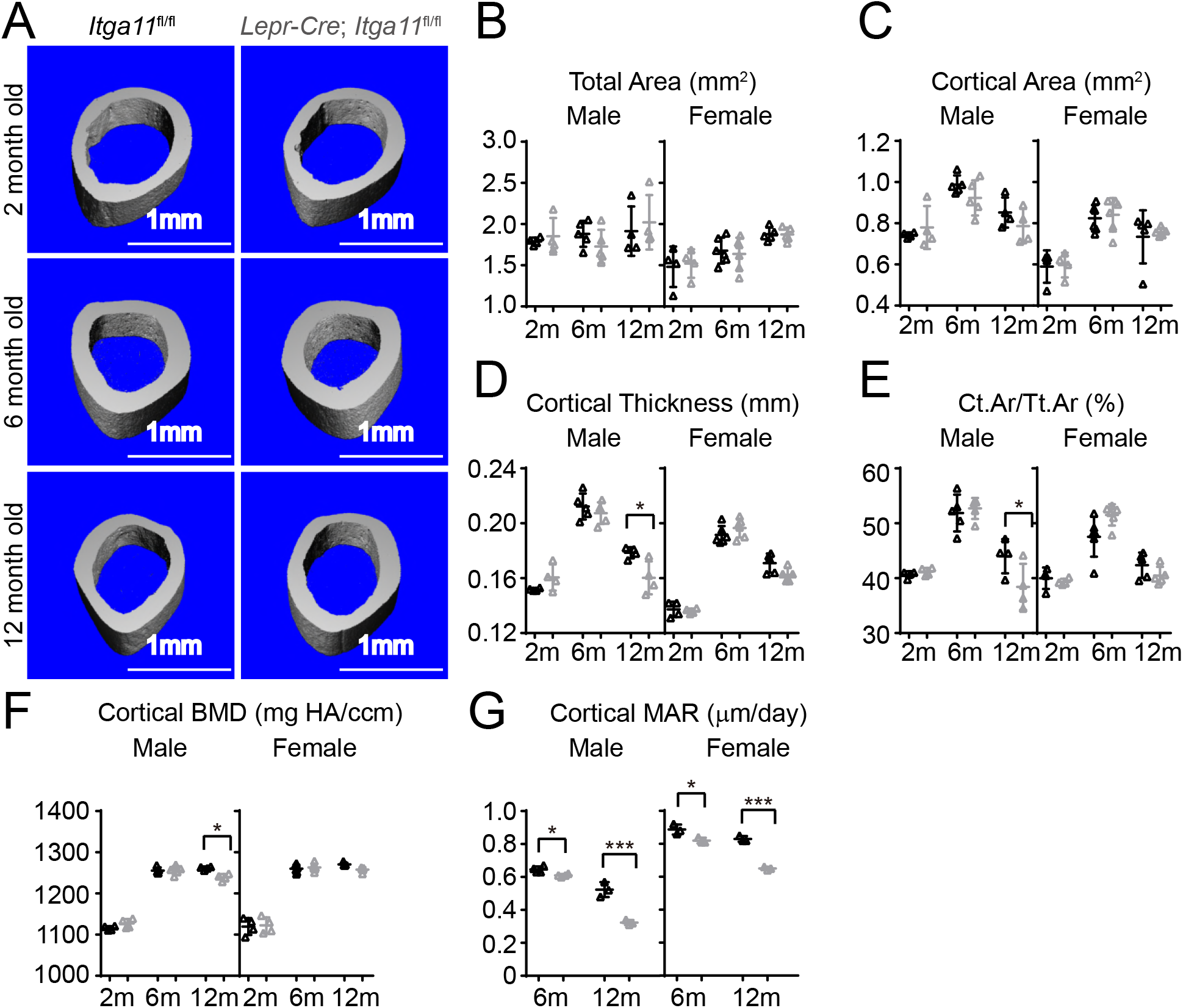
Conditional *Itga11* deletion from LepR^+^ cells reduces cortical bone formation in adult mice. (**A**) Representative microCT images of cortical bone in the mid-femur diaphysis of male *Lepr*-*Cre*; *Itga11^fl/fl^* mice and littermate controls at 2, 6, and 12 months of age. (**B**-**F**) microCT analysis of the total area (**B**), cortical area (**C**), cortical thickness (**D**), cortical area/total area (**E**), and cortical bone mineral density (**F**) in the mid-femur diaphysis of *Lepr*-*Cre*; *Itga11^fl/fl^* mice and sex-matched littermate controls at 2, 6, and 12 months of age (n = 4–6 mice per genotype per sex per time point from at least three independent experiments). (**G**) Cortical bone mineral apposition rate based on calcein double labelling in the mid-femur diaphysis (n = 3–4 mice per genotype per sex per time point from three independent experiments). All numerical data reflect mean ± standard deviation. The statistical significance of differences was determined with two-way ANOVAs with Sidak’s multiple comparisons tests.

### Integrin α11 is required in bone marrow stromal cells to respond to Osteolectin

To test whether Integrin α11 is necessary for the maintenance or the proliferation of skeletal stem/progenitor cells in the bone marrow, we cultured at clonal density enzymatically dissociated femur bone marrow cells from *Lepr*-*Cre*; *Itga11*^fl/fl^ and sex-matched littermate control mice at 2 and 6 months of age. We observed a slight, but statistically significant, reduction in the frequency of cells that formed CFU-F colonies in *Lepr*-*Cre*; *Itga11*^fl/fl^ mice at 2 months of age, though no significant difference was apparent at 6 months of age (Figure 7A). We observed no significant difference in the number of cells per colony at either age (Figure 7B). Integrin α11 is therefore not required for the maintenance of CFU-F *in vivo* or for their proliferation *in vitro*. To test whether integrin α11 regulates the differentiation of bone marrow stromal cells, we cultured CFU-F from *Lepr*-*Cre*; *Itga11*^fl/fl^ and littermate control mice at clonal density, then replated equal numbers of cells from *Itga11* deficient and control colonies into osteogenic or adipogenic culture conditions (Figure 7C and 7D). We also centrifuged 2 × 10^5^ CFU-F cells from *Lepr*-*Cre*; *Itga11*^fl/fl^ and control colonies to form pellets and then cultured them in chondrogenic medium (Figure 7E). Consistent with the decreased osteogenesis from *Itga11* deficient mesenchymal cell lines in culture (Figure 4A) and the reduced osteogenesis in *Lepr*-*Cre*; *Itga11*^fl/fl^ mice *in vivo* (Figure 5 and Figure 6), bone marrow stromal cells from *Lepr*-*Cre*; *Itga11*^fl/fl^ mice formed significantly less bone in culture as compared to control colonies (Figure 7C). This demonstrates that, like Osteolectin, integrin α11 promotes osteogenesis by bone marrow stromal cells. We did not detect any difference between *Lepr*-*Cre*; *Itga11*^fl/fl^ and control colonies in adipogenic or chondrogenic differentiation (Figure 7D and 7E). This is also consistent with the Osteolectin deficiency phenotype, which reduced osteogenesis *in vitro* and *in vivo* without having any detectable effect on adipogenesis or chondrogenesis (Yue et al., 2016).

**Figure 7.**
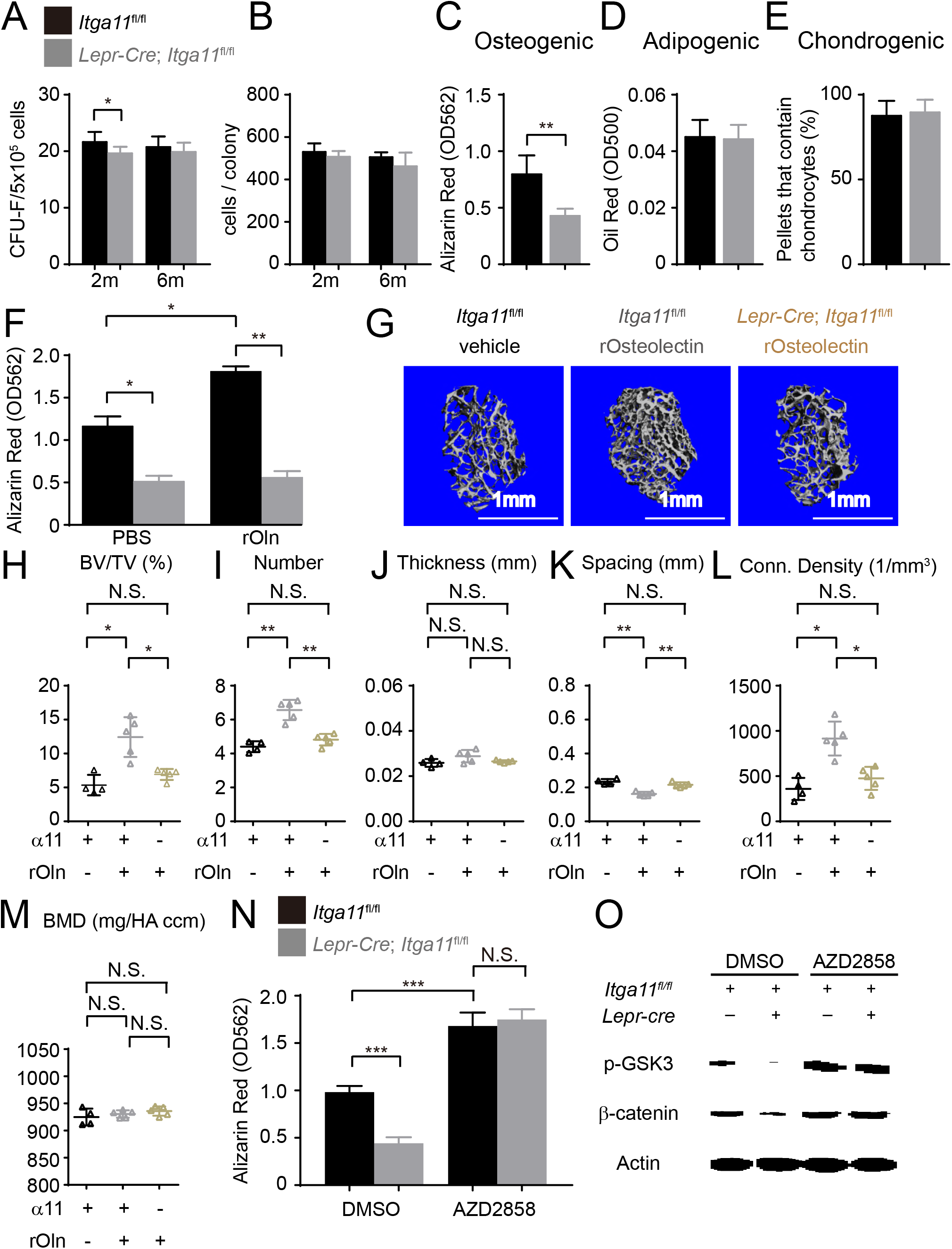
Integrin α11 is required by bone marrow stromal cells to undergo osteogenesis in response to Osteolectin. (**A**) CFU-F frequency and (**B**) cells per CFU-F colony formed by bone marrow cells from *Lepr*-*Cre*; *Itga11^fl/fl^* mice and littermate controls at 2 and 6 months of age (n = 6–8 mice per genotype per time point from at least three independent experiments). (**C-E**) Osteogenic (**C**; n = 7 mice per genotype, total, from seven independent experiments), adipogenic (**D**; n = 6 mice per genotype, total, from six independent experiments), and chondrogenic (**E**; n = 5 mice per genotype, total, from five independent experiments) differentiation of bone marrow stromal cells cultured from the femurs of *Lepr*-*Cre*; *Itga11*^fl/fl^ mice and sex-matched littermate controls at 2 months of age. (**F**) Recombinant mouse Osteolectin promoted osteogenic differentiation in culture by femur bone marrow stromal cells from control but not *Lepr*-*Cre*; *Itga11*^fl/fl^ mice (n = 3 mice per genotype, total, from three independent experiments). (**G**-**M**) Subcutaneous injection of recombinant mouse Osteolectin daily for 28 days (50 µg/kg body mass/day) significantly increased trabecular bone volume and number in female control mice but not female *Lepr*-*Cre*; *Itga11*^fl/fl^ mice. (**G**) Representative microCT images of trabecular bone in the distal femur metaphysis. (**H-M**) microCT analysis of trabecular bone volume/total volume (**H**), trabecular number (**I**), trabecular thickness (**J**), trabecular spacing (**K**), connectivity density (**L**), and bone mineral density (**M**) in the distal femur metaphysis (n = 5 mice per treatment, total, from three independent experiments). (**N**) The GSK3 inhibitor, AZD2858, rescued the osteogenic differentiation in culture of bone marrow stromal cells from the femurs of *Lepr*-*Cre*; *Itga11*^fl/fl^ mice (n = 6 independent experiments in which cells from one mouse of each genotype were cultured in each experiment). (**O**) Western blotting of cultured cell lysates showed that AZD2858 promoted GSK3 phosphorylation and increased β-catenin levels in bone marrow stromal cells from *Lepr*-*Cre*; *Itga11*^fl/fl^ mice and littermate controls. All numerical data reflect mean ± standard deviation. The statistical significance of differences was determined with Wilcoxon’s test followed by Holm-Sidak multiple comparisons adjustment (**A** and **B**), one-way (**F** and **N**) ANOVAs with Sidak’s multiple comparisons tests, by paired t-tests (**C-E**), or by one-way ANOVAs with Tukey’s multiple comparisons tests (**H-M**).

To test whether integrin α11 is necessary for the osteogenic response of bone marrow stromal cells to Osteolectin, we cultured CFU-F from the bone marrow of 2-month-old *Lepr*-*Cre*; *Itga11*^fl/fl^ mice and littermate controls then added osteogenic differentiation medium with or without recombinant mouse Osteolectin. Osteolectin significantly increased osteogenic differentiation by control colonies, but *Lepr*-*Cre*; *Itga11*^fl/fl^ colonies underwent significantly less osteogenesis and did not respond to Osteolectin (Figure 7F). Bone marrow stromal cells thus require integrin α11 to undergo osteogenesis in response to Osteolectin in culture.

To test if bone marrow stromal cells require integrin α11 to undergo osteogenesis in response to Osteolectin *in vivo*, we administered daily subcutaneous injections of recombinant mouse Osteolectin to 2-month-old *Lepr*-*Cre*; *Itga11*^fl/fl^ and littermate control mice for 28 days. Consistent with our prior study (Yue et al., 2016), in the distal femur metaphysis of control mice, Osteolectin treatment significantly increased trabecular bone volume (Figure 7G and 7H), trabecular bone number (Figure 7I), and trabecular connectivity density (Figure 7L), while significantly reducing trabecular spacing (Figure 7K). However, Osteolectin treatment had no significant effect on any of these parameters in *Lepr*-*Cre*; *Itga11*^fl/fl^ mice (Figure 7G-L). LepR^+^ bone marrow stromal cells and their progeny thus require integrin α11 to undergo osteogenesis in response to Osteolectin *in vivo*. Neither Osteolectin administration nor *Itga11* deficiency had any significant effect on cortical bone parameters in this relatively short-term experiment performed in young mice (data not shown).

To test if bone marrow stromal cells from *Lepr*-*Cre*; *Itga11*^fl/fl^ mice retained the capacity to undergo osteogenesis upon Wnt pathway activation, we cultured these cells from 2-month-old *Lepr*-*Cre*; *Itga11*^fl/fl^ and littermate control mice and treated half of the cultures with the Wnt pathway agonist, AZD2858. In control cultures, bone marrow stromal cells from *Lepr*-*Cre*; *Itga11*^fl/fl^ mice underwent significantly less osteogenesis as compared to stromal cells from control mice (Figure 7N). Addition of AZD2858 significantly increased osteogenic differentiation from both *Lepr*-*Cre*; *Itga11*^fl/fl^ and control stromal cells. In cultures treated with DMSO control, *Lepr*-*Cre*; *Itga11*^fl/fl^ stromal cells had lower levels of phosphorylated GSK3 and β-catenin as compared to control stromal cells (Figure 7O). AZD2858 increased the levels of phosphorylated GSK3 and β-catenin in both *Lepr*-*Cre*; *Itga11*^fl/fl^ and control stromal cells (Figure 7O). *Lepr*-*Cre*; *Itga11*^fl/fl^ stromal cells thus retain the ability to undergo osteogenesis in response to Wnt pathway activation, even though they do not respond to Osteolectin.

## Discussion

Our data demonstrate that integrin α11 is a physiologically important receptor for Osteolectin, mediating its effect on osteogenesis. integrin α11 is expressed by LepR^+^ skeletal stem cells and osteoblasts but shows limited expression in non-osteogenic cells (Figure 1G). Osteolectin bound selectively to α11β1 integrin, with nanomolar affinity (Figure 1I and 1J), and promoted Wnt pathway activation in bone marrow stromal cells (Figure 2). integrin α11 was required in bone marrow stromal cells for Wnt pathway activation and osteogenesis in response to Osteolectin (Figure 4A-C). Blocking Wnt pathway activation in bone marrow stromal cells blocked the osteogenic effect of Osteolectin (Figure 3C-D). Conditional deletion of *Itga11* from LepR^+^ cells phenocopied the effect of *Osteolectin* deficiency (Yue et al., 2016): in both cases the mice were grossly normal but exhibited accelerated bone loss during aging, particularly in trabecular bone (Figure 5). Like *Osteolectin* deficiency (Yue et al., 2016), *Itga11* deficiency significantly reduced the rate of bone formation in adult mice (Figure 5M) without affecting the rate of bone resorption (Figure 5N). Bone marrow stromal cells from *Lepr*-*Cre*; *Itga11*^fl/fl^ mice differentiated normally to adipocytes and chondrocytes (Figure 7A-E) but exhibited reduced osteogenic differentiation and did not respond to Osteolectin *in vitro* (Figure 7C and 7F) or *in vivo* (Figure 7G-M). Nonetheless, bone marrow stromal cells from *Lepr*-*Cre*; *Itga11*^fl/fl^ mice retained the ability to form bone in response to a chemical inhibitor of GSK3, which activates the Wnt pathway (Figure 7N and 7O). We conclude that α11 integrin is required by skeletal stem/progenitor cells to undergo osteogenesis in response to Osteolectin.

While deficiency for α11 integrin phenocopied the effects of *Osteolectin* deficiency, we do not rule out a potential role for α10 integrin in mediating certain effects of Osteolectin. α10β1 also bound Osteolectin with nanomolar affinity (Figure 1I). integrin α11 is more highly expressed than a10 by LepR^+^ cells (Figure 1D); however, integrin α10 is expressed by chondrocytes (Bengtsson et al., 2005; Reinisch et al., 2015). While we did not observe any cartilage defects in *Osteolectin* deficient mice (Yue et al., 2016), Osteolectin may promote the differentiation of hypertrophic chondrocytes into bone in adult mice, such as during fracture healing. Therefore, α10 integrin may mediate the effects of Osteolectin on hypertrophic chondrocytes while α11 integrin may mediate the effects of Osteolectin on skeletal stem/progenitor cells.

Osteolectin may not be the only osteogenic ligand for α11 integrin. Collagen is a known ligand for α11β1 integrin (Popova et al., 2007). We found that collagen binds to α11β1 with nanomolar affinity (Figure 1J) but did not detect any effect of exogenous collagen on Wnt pathway activation (Figure 4E) or osteogenic differentiation (Figure 1K). This suggests that collagen may bind α11β1 in a way that regulates cell adhesion and migration but not Wnt pathway activation, at least in skeletal stem/progenitor cells. Alternatively, endogenous collagen may bind α11β1 differently than exogenous collagen, potentially even promoting osteogenesis. Since *Lepr*-*Cre*; *Itga11*^fl/fl^ mice delete *Itga11* in postnatal bone marrow mesenchymal cells that exhibit little contribution to the skeleton during development (Zhou et al., 2014), it remains untested whether α11 integrin regulates osteogenesis during development. If so, this would raise the possibility of a distinct osteogenic ligand for α11 during development as *Osteolectin* deficient mice do not appear to exhibit defects in skeletal development (Yue et al., 2016).

Finally, the osteogenic effect of α11 integrin in mesenchymal cells may not be the only function for α11 integrin. While it is not widely expressed by other cell types, α11 integrin may have other functions in non-mesenchymal cells, or during development, in cells that are not competent to undergo osteogenesis. integrin α11 is expressed by periodontal ligament fibroblasts and is required for the migration of these cells during ligament development, leading to a failure of tooth eruption in germline α11 deficient mice (Popova et al., 2007). This raises the possibility that collagen binding to α11β1 may have biologically distinct consequences in cells that are not competent to form bone.

In conclusion, we identify α11 integrin as an Osteolectin receptor and a new regulator of osteogenesis and adult skeleton maintenance. The identification of a new ligand/receptor pair that regulates the maintenance of the adult skeleton offers the opportunity to better understand the physiological and pathological mechanisms that influence skeletal homeostasis.

## Materials And Methods

### Mice and cell lines

To generate *Itga11*^fl/fl^ mice, CleanCap™ Cas9 mRNA (TriLink) and sgRNAs (transcribed using MEGAshortscript Kit (Ambion), purified using the MEGAclear Kit (Ambion)), and recombineering plasmids were microinjected into C57BL/Ka zygotes. Chimeric mice were genotyped by restriction fragment length poly-morphism (RFLP) analysis and confirmed by Southern blotting and sequencing of the targeted allele. The mice were backcrossed onto a C57BL/Ka background to obtain germline transmission. Mutant mice were backcrossed onto a C57BL/Ka background for at least 3 generations prior to analysis. All procedures were approved by the UTSW Institutional Animal Care and Use Committee (protocol number 2016–101334-G).

Cell lines used in this study included mouse preosteoblast MC3T3-E1 cells (Subclone 4, ATCC CRL-2593), human bone marrow stromal cells from ATCC (PCS-500–012; referred to as hBMSC#1), and human bone marrow stromal cells from Lonza (PT-2501, referred to as hBMSC#2). MC3T3-E1 cells were cultured in Alpha Minimum Essential Medium with ribonucleosides, deoxyribonucleosides, 2 mM L-glutamine and 1 mM sodium pyruvate, but without ascorbic acid (GIBCO, A1049001) supplemented with 10% fetal bovine serum (Sigma, F2442) and penicillin-streptomycin (HyClone). MC3T3-E1 cells were only used for experiments before passage 20. hBMSC cells were cultured in low glucose DMEM (Gibco) supplemented with 20% fetal bovine serum (Sigma, F2442) and penicillin-streptomycin (HyClone), and were only used for experiments before passage 16.

### Western blots

Cells were cultured until confluent, then transferred into osteogenic differentiation medium with or without Osteolectin or small molecule inhibitors of Wnt pathway components. Prior to extracting proteins, cells were washed with PBS and then lysis buffer was added containing 50 mM Tris-HCl, 150 mM NaCl, 1% NP-40, 0.5% sodium deoxycholate, 0.1% SDS, 1 mM sodium vanadate, 0.5 mM sodium fluoride, and cOmplete Mini EDTA-free Protease Inhibitor Cocktail (Sigma). The cells were scraped off the plate in the lysis buffer, transferred to an Eppendorf tube on ice, incubated for 20 minutes with occasional vortexing, then centrifuged at 17,000xg for 10 minutes at 4°C to clear cellular debris. The cell lysates were Western blotted with the indicated antibodies and immunoreactive bands were detected using ECL reagent (Pierce). Antibodies used in this study include anti-Phospho-PI3 Kinase p85(Tyr458)/p55(Tyr199), anti-Phospho-Akt (Ser473), anti-Phospho-GSK-3α/β (Ser21/9), anti-β-catenin, anti-β-Actin, anti-GSK-3β (27C10), and anti-rabbit IgG, HRP-linked antibody from Cell Signaling, anti-mouse Osteolectin (AF3729) and anti-human Osteolectin (AF1904) antibodies from R&D Systems, anti-Adiponectin (ab181699) and anti-integrin α11 antibody (ab198826) from Abcam.

### qRT-PCR

For quantitative reverse transcription PCR (qPCR), cells were lysed using TRIzol LS (Invitrogen). RNA was extracted and reverse transcribed into cDNA using SuperScript III (Invitrogen). qPCR was performed using a Roche LightCycler 480. The primers used for qPCR analysis of mouse RNA include: *Osteolectin*: 5’-AGG TCC TGG GAG GGA GTG-3’ and 5’-GGG CCT CCT GGA GAT TCT T-3’; *Actb*: 5’-GCT CTT TTC CAG CCT TCC TT-3’ and 5’-CTT CTG CAT CCT GTC AGC AA-3’; *Lef1*: 5’-TGT TTA TCC CAT CAC GGG TGG-3’ and 5’-CAT GGA AGT GTC GCC TGA CAG-3’; *Runx2*: 5’-TTA CCT ACA CCC CGC CAG TC-3’ and 5’-TGC TGG TCT GGA AGG GTC C-3’; *Axin2*: 5’-GAG TAG CGC CGT GTT AGT GAC T-3’ and 5’-CCA GGA AAG TCC GGA AGA GGT ATG-3’; *Alp*: 5’-CCA ACT CTT TTG TGC CAG AGA-3’ and 5’-GGC TAC ATT GGT GTT GAG CTT TT-3’. The primers used for qPCR analysis of human RNA include: *Osteolectin*: 5’-ACA TCG TCA CTT ACA TCC TGG GC-3’ and 5’-CAC GCG GGT GTC CAA CG-3’; *Actb*: 5’-ATT GGC AAT GAG CGG TTC-3’ and 5’-CGT GGA TGC CAC AGG ACT-3’; *Lef1*: 5’-TGC CAA ATA TGA ATA ACG ACC CA-3’ and 5’-GAG AAA AGT GCT CGT CAC TGT-3’; *Runx2*: 5’-GAA CCC AGA AGG CAC AGA CA-3’ and 5’-GGC TCA GGT AGG AGG GGT AA-3’; *Axin2*: 5’- CAA CAC CAG GCG GAA CGA A-3’ and 5’- GCC CAA TAA GGA GTG TAA GGA CT-3’; *Alp*: 5’-GTG AAC CGC AAC TGG TAC TC-3’ and 5’-GAG CTG CGT AGC GAT GTC C-3’.

### PCR genotyping

To genotype *Itga11* floxed mice the following primers were used: 5’- AATTCAGTGCCGATCCTCCAGTGTC-3’, 5’-CCCTTGCTTCCTTCTGCTGTCACTT-3’ (*Itga11*^fl^ allele: 370 bp; *Itga11*^+^ allele: 280 bp).

### Integrin binding assay

Integrin binding assays were performed as described (Nishiuchi et al., 2006). Microtiter plates were coated with 10nM recombinant human Osteolectin, recombinant human Pro-Collagen 1 α (R&D Systems), or Bovine Serum Albumin (BSA, Sigma A3156) overnight at 4°C, and then blocked with 10 mg/ml BSA. Recombinant human osteolectin was purified from 293 cells stably expressing Flag-tagged human osteolectin as described (Yue et al., 2016), and 6xHis tagged recombinant human integrin heterodimers were purchased from R&D Systems. The plates were incubated with integrins in TBS buffer (50 mM Tris-Cl, pH 7.5 150 mM NaCl) with 1 mM MnCl_2_, then washed with TBS containing 1 mM MnCl_2_, 0.1% BSA, and 0.02% Tween 20, followed by quantification of bound integrins by an enzyme-linked immunosorbent assay using an anti-His tag monoclonal antibody (Thermo Scientific, clone 4E3D10H2/E3) followed by a horseradish peroxidase-conjugated anti-mouse secondary antibody. After washing, bound HRP was detected using SureBlue TMB Microwell Peroxidase Substrate (KPL) and the reaction was stopped with TMB stop solution (KPL). The optical density was measured at 450 nm.

### MicroCT analysis

MicroCT analysis was performed using the same settings as previously described (Yue et al., 2016). Based on previously described methods (Bouxsein et al., 2010), mouse femurs were dissected, fixed overnight in 4% paraformaldehyde (Thermo Fisher Scientific) and stored in 70% ethanol at 4°C. Femurs and lumbar vertebrae were scanned at an isotropic voxel size of 3.5 mm and 7 mm, respectively, with peak tube voltage of 55 kV and current of 0.145 mA (µCT 35; Scanco). A three-dimensional Gaussian filter (s = 0.8) with a limited, finite filter support of one was used to suppress noise in the images, and a threshold of 263–1000 was used to segment mineralized bone from air and soft tissues. Trabecular bone parameters were measured in the distal metaphysis of the femurs. The region of interest was selected from below the distal growth plate where the epiphyseal cap structure completely disappeared and continued for 100 slices toward the proximal end of the femur. Contours were drawn manually a few voxels away from the endocortical surface to define trabecular bones in the metaphysis. Cortical bone parameters were measured by analyzing 100 slices in mid-diaphysis femurs.

### Statistical analysis

Numbers of experiments noted in figure legends reflect independent experiments performed on different days. Mice were allocated to experiments randomly and samples processed in an arbitrary order, but formal randomization techniques were not used. Prior to analyzing the statistical significance of differences among treatments we tested whether data were normally distributed and whether variance was similar among treatments. To test for normality, we performed the Shapiro–Wilk tests. To test whether variability significantly differed among treatments we performed *F*-tests (for experiments with two treatments) or Levene’s median tests (for experiments with more than two treatments). When the data significantly deviated from normality (*P*<0.01) or variability significantly differed among treatments (*P*<0.05), we log_2_-transformed the data and tested again for normality and variability. If the transformed data no longer significantly deviated from normality and equal variability, we then performed parametric tests on the transformed data. If the transformed data still significantly deviated from normality or equal variability, we performed non-parametric tests on the non-transformed data. Data from the same cell culture experiments were always paired for statistical analysis. Mouse littermates were paired for statistical analysis.

To assess the statistical significance of a difference between two treatments, we used paired two-tailed Student’s *t*-tests (when a parametric test was appropriate) or Wilcoxon’s tests (when a non-parametric test was appropriate). To assess the statistical significance of differences between more than two treatments, we used one-way or two-way repeated measures ANOVAs (when a parametric test was appropriate) followed by post-hoc tests including Dunnett’s, Sidak’s, and Tukey’s tests depending on the experimental settings and planned comparisons, or multiple Wilcoxon’s tests followed by Holm-Sidak’s method for multiple comparisons adjustment (when a non-parametric test was appropriate). Relative mRNA levels were always log2-transformed before any statistical tests were performed. All statistical analyses were performed with Graphpad Prism 7.02. All data represent mean ± standard deviation (*p<0.05, **p<0.01, ***p<0.001).

### Bone marrow digestion and CFU-F assay

As previously described (Yue et al., 2016), mouse femurs and tibias were cut at both ends to flush out intact marrow plugs. Both the flushed plugs and crushed bone metaphyses were subjected to two rounds of enzymatic digestion in prewarmed digestion buffer containing 3 mg/ml type I collagenase (Worthington), 4 mg/ml dispase (Roche Diagnostic) and 1 U/ml DNase I (Sigma) in HBSS with calcium and magnesium, at 37°C for 15 min each round. During each round of digestion, the suspension was vortexed six times for 10 seconds each time at speed level 3 using a Vortex-Genie 2 to promote more complete dissociation. Dissociated cells were transferred into a tube with staining medium (HBSS without calcium and magnesium + 2% fetal bovine serum) and 2 mM EDTA to stop the digestion. Cells were then centrifuged, resuspended in staining medium, and passed through a 90 µm nylon mesh to filter undigested plugs or bone.

To form CFU-F colonies, freshly dissociated bone marrow cell suspensions were plated at clonal density in 6-well plates (5 × 10^5^ cells/well) or 10 cm plates (5 × 10^6^ cells/dish) with DMEM (Gibco) plus 20% fetal bovine serum (Sigma F2442), 10 mM ROCK inhibitor (Y-27632, Selleck), and 1% penicillin/streptomycin (Invitrogen) at 37°C in gas-tight chambers (Billups-Rothenberg) with 1% O_2_ and 6% CO_2_ (with balance Nitrogen) to maintain a low oxygen environment that promoted survival and proliferation (Morrison et al., 2000). The CFU-F culture dish was rinsed with HBBS without calcium and magnesium and replenished with freshly made medium on the second day after plating to wash out contaminating macrophages. Cultures were then maintained in a gas-tight chamber that was flushed daily for 1 minute with a custom low oxygen gas mixture (1% O_2_, 6% CO_2_, balance Nitrogen). The culture medium was changed every 4 days. To count CFU-F colonies, the cultures were stained with 0.1% Toluidine blue in 4% formalin solution eight days after plating.

### *in vitro* differentiation assay

The osteogenic potential of primary CFU-F cells, human bone marrow stromal cells, and MC3T3-E1 cells was assessed by plating the cells into 48- well plates (25,000 cells/cm^2^). On the second day after plating, the culture medium was replaced with osteogenic differentiation medium (StemPro Osteogenesis Differentiation kit, Gibco). Cells were maintained in the differentiation medium, with medium change every other day for 14 days for primary CFU-F cells and MC3T3-E1 cells before differentiation was assessed. For human bone marrow stromal cells, the culture medium was changed every 3 days for 21 days. Osteoblastic differentiation was detected by staining with Alizarin red S (Sigma). To quantitate Alizarin red staining, the stained cells were rinsed with PBS, and extracted with 10% (w/v) cetylpyridinium chloride in 10 mM sodium phosphate, pH 7.0 for 10 minutes at room temperature. Alizarin red in the extract was quantitated by optical density measurement at 562 nm.

The adipogenic potential of CFU-F cells was assessed by plating them into 48-well plates (25,000 cells/cm^2^). On the second day after plating, the culture medium was replaced with adipogenic differentiation medium (StemPro Adipogenesis Differentiation Kit, Gibco) and the cultures were allowed to differentiate for 14 days, with culture medium changed every 3 days. Adipocyte differentiation was detected by staining with Oil red O (Sigma). To quantitate the amount of Oil red O staining, cells were rinsed with PBS, and extracted with 100% isopropanol for 10 minutes at room temperature. Oil red O in the extract was quantitated by optical density measurement at 500 nm.

The chondrogenic potential of CFU-F cells was assessed by centrifuging 2 × 10^5^ cells to form cell pellets, which were then cultured in chondrogenic medium (StemPro chondrogenesis differentiation kit; Gibco) for 21 days. The culture medium was changed every 3 days. Chondrocyte formation within the cell pellets was assessed by cryosectioning and Toluidine blue staining as described (Robey et al., 2014).

### Calcein double labeling and histomorphometry analysis

As previously described (Egan et al., 2012), mice were injected intraperitoneally with 10mg/kg body weight of calcein, dissolved in 0.15 M NaCl plus 2% NaHCO_3_ in water, at day 0 and day 7. Mice were sacrificed on day 9. Mouse tibias were fixed overnight in 4% paraformaldehyde at 4°C, dehydrated in 30% sucrose at 4°C for two days and sectioned without decalcification (7 µm sections). Mineral apposition rates were determined as previously described (Egan et al., 2012).

### ELISA assays

The bone resorption rate was determined by measuring urinary levels of deoxypyridinoline (DPD) using a MicroVue DPD ELISA Kit (Quidel). The DPD values were normalized to urinary creatinine levels using the MicroVue Creatinine Assay Kit (Quidel). The ELISA assay for Osteolectin was described previously (Yue et al., 2016).

## Acknowledgements

SJM is a Howard Hughes Medical Institute (HHMI) Investigator, the Mary McDermott Cook Chair in Pediatric Genetics, the Kathryn and Gene Bishop Distinguished Chair in Pediatric Research, the director of the Hamon Laboratory for Stem Cells and Cancer, and a Cancer Prevention and Research Institute of Texas Scholar. BS was supported by a Ruth L. Kirschstein National Research Service Award (NRSA) Postdoctoral Fellowship (F32) from the National Heart, Lung, and Blood Institute (1F32HL139016–01). AT was supported by the Leopoldina Fellowship Program (LPDS 2016–16) from the German National Academy of Sciences. We thank Nicolas Loof and the Moody Foundation Flow Cytometry Facility, Albert Gross for mouse colony management, Yu Zhang and Hao Zhu for intracytoplasmic sperm injection. This work was supported by the National Institute on Aging (R37 AG02494514).

## Figure Legends

**Table S1.** Markers used to isolate bone marrow cell populations. Stem and progenitor cell populations were isolated from mouse bone marrow by flow cytometry using the listed markers. Lineage (Lin) markers used to isolate Lineage negative cell populations were CD2, CD3, CD5, CD8, Ter119, Gr-1, and B220.

| Full description | Abbreviation | Marker Used to Isolate | References |
| --- | --- | --- | --- |
| Bone Marrow Cells | BMC | Ter119<sup>-</sup> |  |
| Endothelial Cells | Endo | VE-cadherin<sup>+</sup>, CD45<sup>-</sup>, Ter119<sup>-</sup> |  |
| LepR<sup>+</sup> Cells | LepR<sup>+</sup> | LepR<sup>+</sup>, CD45<sup>-</sup>, Ter119<sup>-</sup>, CD31<sup>-</sup> | (Zhou et al., 2014) |
| Osteoblasts | OB | *Col2.3-GFP*<sup>+</sup> | (Kalajzic et al., 2002) |
| Common Myeloid Progenitors | CMP | Lin<sup>-</sup>, c-kit<sup>+</sup>, Sca1<sup>-</sup>, CD34<sup>+</sup>, CD16/32<sup>-</sup> | (Akashi et al., 2000) |
| Granulocyte and Monocyte Precursors | GMP | Lin<sup>-</sup>, c-kit<sup>+</sup>, Sca1<sup>-</sup>, CD34<sup>+</sup>, CD16/32<sup>+</sup> | (Akashi et al., 2000) |
| Megakaryocyte and Erythroid precursors | MEP | Lin<sup>-</sup>, c-kit<sup>+</sup>, Sca1<sup>-</sup>, CD34<sup>-</sup>, CD16/32<sup>-</sup> | (Akashi et al., 2000) |
| Common Lymphoid Progenitors | CLP | Lin<sup>-</sup>, c-kit<sup>low</sup>, Sca1<sup>low</sup>, CD127<sup>+</sup>, CD135<sup>+</sup> | (Kondo et al., 1997) |
| CD3<sup>+</sup> T Cells | CD3<sup>+</sup> | Ter119<sup>-</sup>, CD3<sup>+</sup> |  |
| Erythroid cells | Erythroid | Ter119<sup>+</sup>, CD71<sup>+</sup> |  |
| Myeloid cells | Myeloid | Mac-1<sup>+</sup>, Gr-1<sup>+</sup> |  |
| c-kit<sup>+</sup> Progenitors | LSK | Lin<sup>-</sup>, c-kit<sup>+</sup>, Sca1<sup>+</sup> | (Okada et al., 1992) |
| Multipotent Progenitors | MPP | Lin<sup>-</sup>, c-kit<sup>+</sup>, Sca1<sup>+</sup>, CD150<sup>-</sup>, CD48<sup>-</sup> | (Kiel et al., 2008) |
| Hematopoietic Stem Cells | HSC | Lin<sup>-</sup>, c-kit<sup>+</sup>, Sca1<sup>+</sup>, CD150<sup>+</sup>, CD48<sup>-</sup> | (Kiel et al., 2005) |
| B220<sup>+</sup> B Cells | B220<sup>+</sup> | Ter119<sup>-</sup>, B220<sup>+</sup> |  |
| Pro-B Cells | Pro-B | B220<sup>+</sup>, CD43<sup>+</sup>, CD24<sup>+</sup>, IgM<sup>-</sup> | (Li et al., 1996) |
| Pre-pro-B Cells | Prepro-B | B220<sup>+</sup>, CD43<sup>+</sup>, CD24<sup>-</sup>, IgM<sup>-</sup> | (Li et al., 1996) |

**Figure S5A.**
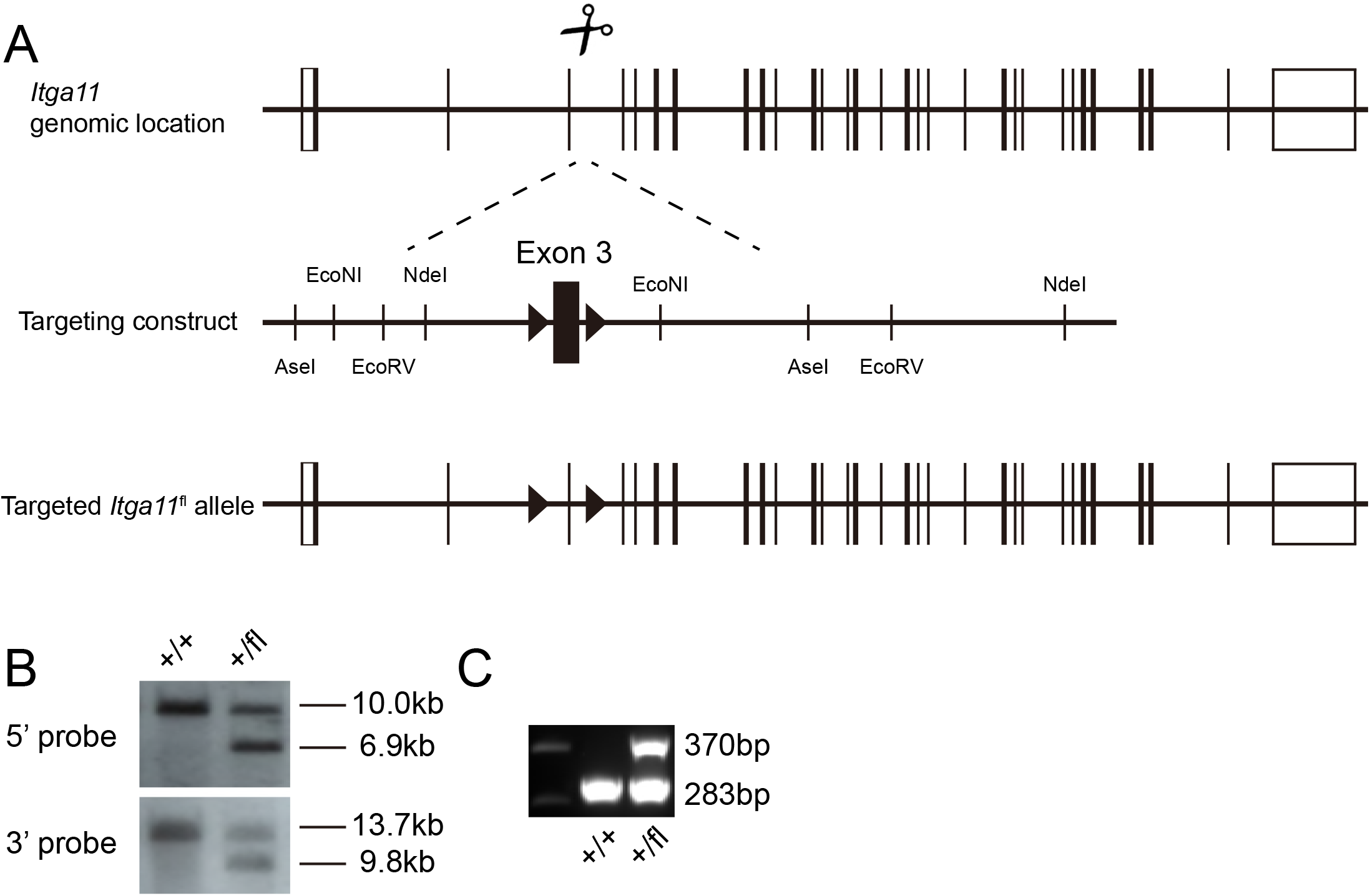
Generation of an *Itga11* floxed mice. (**A**) DNA surrounding *Itga11* exon3 was used as homologous arms for recombineering. The recombineering plasmid was modified by inserting loxP sites flanking exon3. Deletion of exon 3 leads to a frame shift that would be expected to give a complete loss of α11 function (Popova et al., 2007). The loxP insertion sites were chosen so as not to disrupt sequences conserved among species. Cas9 mRNA, sgRNAs, and the recombineering plasmid were microinjected into C57BL/Ka zygotes. Correctly-targeted founder mice (F0) were identified by Southern blotting (**B**) using an internal (3’) probe and an external (5’) probe to ensure the recombineering vector was integrated in the correct genomic location, and not in other regions of the genome. Mice were backcrossed at least 3 times onto a C57BL/Ka background before analysis. (**C**) PCR genotyping demonstrated germline transmission of the *Itga11*^flox^ allele.

## References

Ai, M., S.L. Holmen, W. Van Hul, B.O. Williams, and M.L. Warman. 2005. Reduced affinity to and inhibition by DKK1 form a common mechanism by which high bone mass-associated missense mutations in LRP5 affect canonical Wnt signaling. Mol Cell Biol. 25:4946–4955.

Akashi, K., D. Traver, T. Miyamoto, and I.L. Weissman. 2000. A clonogenic common myeloid progenitor that gives rise to all myeloid lineages. Nature. 404:193–197.

Aszodi, A., E.B. Hunziker, C. Brakebusch, and R. Fassler. 2003. Beta1 integrins regulate chondrocyte rotation, G1 progression, and cytokinesis. Genes Dev. 17:2465–2479.

Balemans, W., M. Ebeling, N. Patel, E. Van Hul, P. Olson, M. Dioszegi, C. Lacza, W. Wuyts, J. Van Den Ende, P. Willems, A.F. Paes-Alves, S. Hill, M. Bueno, F.J. Ramos, P. Tacconi, F.G. Dikkers, C. Stratakis, K. Lindpaintner, B. Vickery, D. Foernzler, and W. Van Hul. 2001. Increased bone density in sclerosteosis is due to the deficiency of a novel secreted protein (SOST). Hum Mol Genet. 10:537–543.

Bannwarth, S., V. Giordanengo, J. Grosgeorge, C. Turc-Carel, and J.C. Lefebvre. 1999. Cloning, mapping, and genomic organization of the LSLCL gene, encoding a new lymphocytic secreted mucin-like protein with a C-type lectin domain: A new model of exon shuffling. Genomics. 57:316–317.

Bannwarth, S., V. Giordanengo, J. Lesimple, and J.C. Lefebvre. 1998. Molecular cloning of a new secreted sulfated mucin-like protein with a C-type lectin domain that is expressed in lymphoblastic cells. J Biol Chem. 273:1911–1916.

Bengtsson, T., A. Aszodi, C. Nicolae, E.B. Hunziker, E. Lundgren-Akerlund, and R. Fassler. 2005. Loss of alpha10beta1 integrin expression leads to moderate dysfunction of growth plate chondrocytes. J Cell Sci. 118:929–936.

Bennett, C.N., K.A. Longo, W.S. Wright, L.J. Suva, T.F. Lane, K.D. Hankenson, and O.A. MacDougald. 2005. Regulation of osteoblastogenesis and bone mass by Wnt10b. Proc Natl Acad Sci U S A. 102:3324–3329.

Berg, S., M. Bergh, S. Hellberg, K. Hogdin, Y. Lo-Alfredsson, P. Soderman, S. von Berg, T. Weigelt, M. Ormo, Y. Xue, J. Tucker, J. Neelissen, E. Jerning, Y. Nilsson, and R. Bhat. 2012. Discovery of novel potent and highly selective glycogen synthase kinase-3beta (GSK3beta) inhibitors for Alzheimer’s disease: design, synthesis, and characterization of pyrazines. J Med Chem. 55:9107–9119.

Bouxsein, M.L., S.K. Boyd, B.A. Christiansen, R.E. Guldberg, K.J. Jepsen, and R. Muller. 2010. Guidelines for assessment of bone microstructure in rodents using micro-computed tomography. J Bone Miner Res. 25:1468–1486.

Boyden, L.M., J. Mao, J. Belsky, L. Mitzner, A. Farhi, M.A. Mitnick, D. Wu, K. Insogna, and R.P. Lifton. 2002. High bone density due to a mutation in LDL-receptor-related protein 5. N Engl J Med. 346:1513–1521.

Burkhalter, R.J., J. Symowicz, L.G. Hudson, C.J. Gottardi, and M.S. Stack. 2011. Integrin regulation of beta-catenin signaling in ovarian carcinoma. J Biol Chem. 286:23467–23475.

Chan, C.K., E.Y. Seo, J.Y. Chen, D. Lo, A. McArdle, R. Sinha, R. Tevlin, J. Seita, J. Vincent-Tompkins, T. Wearda, W.J. Lu, K. Senarath-Yapa, M.T. Chung, O. Marecic, M. Tran, K.S. Yan, R. Upton, G.G. Walmsley, A.S. Lee, D. Sahoo, C.J. Kuo, I.L. Weissman, and M.T. Longaker. 2015. Identification and specification of the mouse skeletal stem cell. Cell. 160:285–298.

Chen, B., M.E. Dodge, W. Tang, J. Lu, Z. Ma, C.W. Fan, S. Wei, W. Hao, J. Kilgore, N.S. Williams, M.G. Roth, J.F. Amatruda, C. Chen, and L. Lum. 2009. Small molecule-mediated disruption of Wnt-dependent signaling in tissue regeneration and cancer. Nat Chem Biol. 5:100–107.

Chen, Q., P. Shou, L. Zhang, C. Xu, C. Zheng, Y. Han, W. Li, Y. Huang, X. Zhang, C. Shao, A.I. Roberts, A.B. Rabson, G. Ren, Y. Zhang, Y. Wang, D.T. Denhardt, and Y. Shi. 2014. An osteopontin-integrin interaction plays a critical role in directing adipogenesis and osteogenesis by mesenchymal stem cells. Stem Cells. 32:327–337.

Cross, D.A., D.R. Alessi, P. Cohen, M. Andjelkovich, and B.A. Hemmings. 1995. Inhibition of glycogen synthase kinase-3 by insulin mediated by protein kinase B. Nature. 378:785–789.

Dejaeger, M., A.M. Bohm, N. Dirckx, J. Devriese, E. Nefyodova, R. Cardoen, R. St-Arnaud, J. Tournoy, F.P. Luyten, and C. Maes. 2017. Integrin-Linked Kinase Regulates Bone Formation by Controlling Cytoskeletal Organization and Modulating BMP and Wnt Signaling in Osteoprogenitors. J Bone Miner Res. 32:2087–2102.

Delcommenne, M., C. Tan, V. Gray, L. Rue, J. Woodgett, and S. Dedhar. 1998. Phosphoinositide-3-OH kinase-dependent regulation of glycogen synthase kinase 3 and protein kinase B/AKT by the integrin-linked kinase. Proc Natl Acad Sci U S A. 95:11211–11216.

Ding, L., and S.J. Morrison. 2013. Haematopoietic stem cells and early lymphoid progenitors occupy distinct bone marrow niches. Nature. 495:231–235.

Ding, L., T.L. Saunders, G. Enikolopov, and S.J. Morrison. 2012. Endothelial and perivascular cells maintain haematopoietic stem cells. Nature. 481:457–462.

Dong, Y.F., Y. Soung do, E.M. Schwarz, R.J. O’Keefe, and H. Drissi. 2006. Wnt induction of chondrocyte hypertrophy through the Runx2 transcription factor. J Cell Physiol. 208:77–86.

Dy, P., W. Wang, P. Bhattaram, Q. Wang, L. Wang, R.T. Ballock, and V. Lefebvre. 2012. Sox9 directs hypertrophic maturation and blocks osteoblast differentiation of growth plate chondrocytes. Dev Cell. 22:597–609.

Egan, K.P., T.A. Brennan, and R.J. Pignolo. 2012. Bone histomorphometry using free and commonly available software. Histopathology. 61:1168–1173.

Engel, B.E., E. Welsh, M.F. Emmons, P.G. Santiago-Cardona, and W.D. Cress. 2013. Expression of integrin alpha 10 is transcriptionally activated by pRb in mouse osteoblasts and is downregulated in multiple solid tumors. Cell Death Dis. 4:e938.

Filali, M., N. Cheng, D. Abbott, V. Leontiev, and J.F. Engelhardt. 2002. Wnt-3A/beta-catenin signaling induces transcription from the LEF-1 promoter. J Biol Chem. 277:33398–33410.

Fong, S., S. Jones, M.E. Renz, H.H. Chiu, A.M. Ryan, L.G. Presta, and D. Jackson. 1997. Mucosal addressin cell adhesion molecule-1 (MAdCAM-1). Its binding motif for alpha 4 beta 7 and role in experimental colitis. Immunol Res. 16:299–311.

Gardner, J.M., and R.O. Hynes. 1985. Interaction of fibronectin with its receptor on platelets. Cell. 42:439–448.

Gaur, T., C.J. Lengner, H. Hovhannisyan, R.A. Bhat, P.V. Bodine, B.S. Komm, A. Javed, A.J. van Wijnen, J.L. Stein, G.S. Stein, and J.B. Lian. 2005. Canonical WNT signaling promotes osteogenesis by directly stimulating Runx2 gene expression. J Biol Chem. 280:33132–33140.

Gilmour, P.S., P.J. O’Shea, M. Fagura, J.E. Pilling, H. Sanganee, H. Wada, P.F. Courtney, S. Kavanagh, P.A. Hall, and K.J. Escott. 2013. Human stem cell osteoblastogenesis mediated by novel glycogen synthase kinase 3 inhibitors induces bone formation and a unique bone turnover biomarker profile in rats. Toxicol Appl Pharmacol. 272:399–407.

Gong, Y., R.B. Slee, N. Fukai, G. Rawadi, S. Roman-Roman, A.M. Reginato, H. Wang, T. Cundy, F.H. Glorieux, D. Lev, M. Zacharin, K. Oexle, J. Marcelino, W. Suwairi, S. Heeger, G. Sabatakos, S. Apte, W.N. Adkins, J. Allgrove, M. Arslan-Kirchner, J.A. Batch, P. Beighton, G.C. Black, R.G. Boles, L.M. Boon, C. Borrone, H.G. Brunner, G.F. Carle, B. Dallapiccola, A. De Paepe, B. Floege, M.L. Halfhide, B. Hall, R.C. Hennekam, T. Hirose, A. Jans, H. Juppner, C.A. Kim, K. Keppler-Noreuil, A. Kohlschuetter, D. LaCombe, M. Lambert, E. Lemyre, T. Letteboer, L. Peltonen, R.S. Ramesar, M. Romanengo, H. Somer, E. Steichen-Gersdorf, B. Steinmann, B. Sullivan, A. Superti-Furga, W. Swoboda, M.J. van den Boogaard, W. Van Hul, M. Vikkula, M. Votruba, B. Zabel, T. Garcia, R. Baron, B.R. Olsen, M.L. Warman, and G. Osteoporosis-Pseudoglioma Syndrome Collaborative. 2001. LDL receptor-related protein 5 (LRP5) affects bone accrual and eye development. Cell. 107:513–523.

Hamidouche, Z., O. Fromigue, J. Ringe, T. Haupl, P. Vaudin, J.C. Pages, S. Srouji, E. Livne, and P.J. Marie. 2009. Priming integrin alpha5 promotes human mesenchymal stromal cell osteoblast differentiation and osteogenesis. Proc Natl Acad Sci U S A. 106:18587–18591.

Hernandez, L., K.J. Roux, E.S. Wong, L.C. Mounkes, R. Mutalif, R. Navasankari, B. Rai, S. Cool, J.W. Jeong, H. Wang, H.S. Lee, S. Kozlov, M. Grunert, T. Keeble, C.M. Jones, M.D. Meta, S.G. Young, I.O. Daar, B. Burke, A.O. Perantoni, and C.L. Stewart. 2010. Functional coupling between the extracellular matrix and nuclear lamina by Wnt signaling in progeria. Dev Cell. 19:413–425.

Himburg, H.A., C.M. Termini, L. Schlussel, J. Kan, M. Li, L. Zhao, T. Fang, J.P. Sasine, V.Y. Chang, and J.P. Chute. 2018. Distinct Bone Marrow Sources of Pleiotrophin Control Hematopoietic Stem Cell Maintenance and Regeneration. Cell Stem Cell.

Hiraoka, A., A. Sugimura, T. Seki, T. Nagasawa, N. Ohta, M. Shimonishi, M. Hagiya, and S. Shimizu. 1997. Cloning, expression, and characterization of a cDNA encoding a novel human growth factor for primitive hematopoietic progenitor cells. Proc Natl Acad Sci U S A. 94:7577–7582.

Hiraoka, A., K. Yano Ki, N. Kagami, K. Takeshige, H. Mio, H. Anazawa, and S. Sugimoto. 2001. Stem cell growth factor: in situ hybridization analysis on the gene expression, molecular characterization and in vitro proliferative activity of a recombinant preparation on primitive hematopoietic progenitor cells. Hematol J. 2:307–315.

Holmen, S.L., T.A. Giambernardi, C.R. Zylstra, B.D. Buckner-Berghuis, J.H. Resau, J.F. Hess, V. Glatt, M.L. Bouxsein, M. Ai, M.L. Warman, and B.O. Williams. 2004. Decreased BMD and limb deformities in mice carrying mutations in both Lrp5 and Lrp6. J Bone Miner Res. 19:2033–2040.

Hovanes, K., T.W. Li, J.E. Munguia, T. Truong, T. Milovanovic, J. Lawrence Marsh, R.F. Holcombe, and M.L. Waterman. 2001. Beta-catenin-sensitive isoforms of lymphoid enhancer factor-1 are selectively expressed in colon cancer. Nat Genet. 28:53–57.

Humphries, J.D., A. Byron, and M.J. Humphries. 2006. Integrin ligands at a glance. J Cell Sci. 119:3901–3903.

Hynes, R.O. 1992. Integrins: versatility, modulation, and signaling in cell adhesion. Cell. 69:11–25.

Jho, E.H., T. Zhang, C. Domon, C.K. Joo, J.N. Freund, and F. Costantini. 2002. Wnt/beta-catenin/Tcf signaling induces the transcription of Axin2, a negative regulator of the signaling pathway. Mol Cell Biol. 22:1172–1183.

Kalajzic, Z., P. Liu, I. Kalajzic, Z. Du, A. Braut, M. Mina, E. Canalis, and D.W. Rowe. 2002. Directing the expression of a green fluorescent protein transgene in differentiated osteoblasts: comparison between rat type I collagen and rat osteocalcin promoters. Bone. 31:654–660.

Kaltz, N., J. Ringe, C. Holzwarth, P. Charbord, M. Niemeyer, V.R. Jacobs, C. Peschel, T. Haupl, and R.A. Oostendorp. 2010. Novel markers of mesenchymal stem cells defined by genome-wide gene expression analysis of stromal cells from different sources. Exp Cell Res. 316:2609–2617.

Karsenty, G. 2003. The complexities of skeletal biology. Nature. 423:316–318.

Kato, M., M.S. Patel, R. Levasseur, I. Lobov, B.H. Chang, D.A. Glass, 2nd, C. Hartmann, L. Li, T.H. Hwang, C.F. Brayton, R.A. Lang, G. Karsenty, and L. Chan. 2002. Cbfa1-independent decrease in osteoblast proliferation, osteopenia, and persistent embryonic eye vascularization in mice deficient in Lrp5, a Wnt coreceptor. J Cell Biol. 157:303–314.

Kiel, M.J., O.H. Yilmaz, T. Iwashita, O.H. Yilmaz, C. Terhorst, and S.J. Morrison. 2005. SLAM family receptors distinguish hematopoietic stem and progenitor cells and reveal endothelial niches for stem cells. Cell. 121:1109–1121.

Kiel, M.J., O.H. Yilmaz, and S.J. Morrison. 2008. CD150-cells are transiently reconstituting multipotent progenitors with little or no stem cell activity. Blood. 111:4413–4414.

Kondo, M., I.L. Weissman, and K. Akashi. 1997. Identification of clonogenic common lymphoid progenitors in mouse bone marrow. Cell. 91:661–672.

Krishnan, V., H.U. Bryant, and O.A. Macdougald. 2006. Regulation of bone mass by Wnt signaling. J Clin Invest. 116:1202–1209.

Kronenberg, H.M. 2003. Developmental regulation of the growth plate. Nature. 423:332–336.

Kulkarni, N.H., J.E. Onyia, Q. Zeng, X. Tian, M. Liu, D.L. Halladay, C.A. Frolik, T. Engler, T. Wei, A. Kriauciunas, T.J. Martin, M. Sato, H.U. Bryant, and Y.L. Ma. 2006. Orally bioavailable GSK-3alpha/beta dual inhibitor increases markers of cellular differentiation in vitro and bone mass in vivo. J Bone Miner Res. 21:910–920.

Li, Y.S., R. Wasserman, K. Hayakawa, and R.R. Hardy. 1996. Identification of the earliest B lineage stage in mouse bone marrow. Immunity. 5:527–535.

Lustig, B., B. Jerchow, M. Sachs, S. Weiler, T. Pietsch, U. Karsten, M. van de Wetering, H. Clevers, P.M. Schlag, W. Birchmeier, and J. Behrens. 2002. Negative feedback loop of Wnt signaling through upregulation of conductin/axin2 in colorectal and liver tumors. Mol Cell Biol. 22:1184–1193.

Marsell, R., G. Sisask, Y. Nilsson, A.K. Sundgren-Andersson, U. Andersson, S. Larsson, O. Nilsson, O. Ljunggren, and K.B. Jonsson. 2012. GSK-3 inhibition by an orally active small molecule increases bone mass in rats. Bone. 50:619–627.

Mizoguchi, T., S. Pinho, J. Ahmed, Y. Kunisaki, M. Hanoun, A. Mendelson, N. Ono, H.M. Kronenberg, and P.S. Frenette. 2014. Osterix marks distinct waves of primitive and definitive stromal progenitors during bone marrow development. Dev Cell. 29:340–349.

Morrison, S.J., S.E. Perez, Z. Qiao, J.M. Verdi, C. Hicks, G. Weinmaster, and D.J. Anderson. 2000. Transient Notch activation initiates an irreversible switch from neurogenesis to gliogenesis by neural crest stem cells. Cell. 101:499–510.

Moursi, A.M., R.K. Globus, and C.H. Damsky. 1997. Interactions between integrin receptors and fibronectin are required for calvarial osteoblast differentiation in vitro. J Cell Sci. 110 (Pt 18):2187–2196.

Nishiuchi, R., J. Takagi, M. Hayashi, H. Ido, Y. Yagi, N. Sanzen, T. Tsuji, M. Yamada, and K. Sekiguchi. 2006. Ligand-binding specificities of laminin-binding integrins: a comprehensive survey of laminin-integrin interactions using recombinant alpha3beta1, alpha6beta1, alpha7beta1 and alpha6beta4 integrins. Matrix Biol. 25:189–197.

Novak, A., S.C. Hsu, C. Leung-Hagesteijn, G. Radeva, J. Papkoff, R. Montesano, C. Roskelley, R. Grosschedl, and S. Dedhar. 1998. Cell adhesion and the integrin-linked kinase regulate the LEF-1 and beta-catenin signaling pathways. Proc Natl Acad Sci U S A. 95:4374–4379.

Oguro, H., L. Ding, and S.J. Morrison. 2013. SLAM family markers resolve functionally distinct subpopulations of hematopoietic stem cells and multipotent progenitors. Cell Stem Cell. 13:102–116.

Okada, S., H. Nakauchi, K. Nagayoshi, S. Nishikawa, Y. Miura, and T. Suda. 1992. In vivo and in vitro stem cell function of c-kit-and Sca-1-positive murine hematopoietic cells. Blood. 80:3044–3050.

Peifer, M., L.M. Pai, and M. Casey. 1994. Phosphorylation of the Drosophila adherens junction protein Armadillo: roles for wingless signal and zeste-white 3 kinase. Dev Biol. 166:543–556.

Pierschbacher, M.D., and E. Ruoslahti. 1984. Cell attachment activity of fibronectin can be duplicated by small synthetic fragments of the molecule. Nature. 309:30–33.

Plow, E.F., M.D. Pierschbacher, E. Ruoslahti, G.A. Marguerie, and M.H. Ginsberg. 1985. The effect of Arg-Gly-Asp-containing peptides on fibrinogen and von Willebrand factor binding to platelets. Proc Natl Acad Sci U S A. 82:8057–8061.

Popova, S.N., M. Barczyk, C.F. Tiger, W. Beertsen, P. Zigrino, A. Aszodi, N. Miosge, E. Forsberg, and D. Gullberg. 2007. Alpha11 beta1 integrin-dependent regulation of periodontal ligament function in the erupting mouse incisor. Mol Cell Biol. 27:4306–4316.

Popova, S.N., B. Rodriguez-Sanchez, A. Liden, C. Betsholtz, T. Van Den Bos, and D. Gullberg. 2004. The mesenchymal alpha11beta1 integrin attenuates PDGF-BB-stimulated chemotaxis of embryonic fibroblasts on collagens. Dev Biol. 270:427–442.

Raducanu, A., E.B. Hunziker, I. Drosse, and A. Aszodi. 2009. Beta1 integrin deficiency results in multiple abnormalities of the knee joint. J Biol Chem. 284:23780–23792.

Rallis, C., S.M. Pinchin, and D. Ish-Horowicz. 2010. Cell-autonomous integrin control of Wnt and Notch signalling during somitogenesis. Development. 137:3591–3601.

Rawadi, G., B. Vayssiere, F. Dunn, R. Baron, and S. Roman-Roman. 2003. BMP-2 controls alkaline phosphatase expression and osteoblast mineralization by a Wnt autocrine loop. J Bone Miner Res. 18:1842–1853.

Reinisch, A., N. Etchart, D. Thomas, N.A. Hofmann, M. Fruehwirth, S. Sinha, C.K. Chan, K. Senarath-Yapa, E.Y. Seo, T. Wearda, U.F. Hartwig, C. Beham-Schmid, S. Trajanoski, Q. Lin, W. Wagner, C. Dullin, F. Alves, M. Andreeff, I.L. Weissman, M.T. Longaker, K. Schallmoser, R. Majeti, and D. Strunk. 2015. Epigenetic and in vivo comparison of diverse MSC sources reveals an endochondral signature for human hematopoietic niche formation. Blood. 125:249–260.

Robey, P.G., S.A. Kuznetsov, M. Riminucci, and P. Bianco. 2014. Bone marrow stromal cell assays: in vitro and in vivo. Methods Mol Biol. 1130:279–293.

Rodda, S.J., and A.P. McMahon. 2006. Distinct roles for Hedgehog and canonical Wnt signaling in specification, differentiation and maintenance of osteoblast progenitors. Development. 133:3231–3244.

Shekaran, A., J.T. Shoemaker, T.E. Kavanaugh, A.S. Lin, M.C. LaPlaca, Y. Fan, R.E. Guldberg, and A.J. Garcia. 2014. The effect of conditional inactivation of beta 1 integrins using twist 2 Cre, Osterix Cre and osteocalcin Cre lines on skeletal phenotype. Bone. 68:131–141.

Sisask, G., R. Marsell, A. Sundgren-Andersson, S. Larsson, O. Nilsson, O. Ljunggren, and K.B. Jonsson. 2013. Rats treated with AZD2858, a GSK3 inhibitor, heal fractures rapidly without endochondral bone formation. Bone. 54:126–132.

Sun, C., H. Yuan, L. Wang, X. Wei, L. Williams, P.H. Krebsbach, J.L. Guan, and F. Liu. 2016. FAK Promotes Osteoblast Progenitor Cell Proliferation and Differentiation by Enhancing Wnt Signaling. J Bone Miner Res. 31:2227–2238.

Velling, T., M. Kusche-Gullberg, T. Sejersen, and D. Gullberg. 1999. cDNA cloning and chromosomal localization of human alpha (11) integrin. A collagen-binding, I domain-containing, beta (1)-associated integrin alpha-chain present in muscle tissues. J Biol Chem. 274:25735–25742.

Viney, J.L., S. Jones, H.H. Chiu, B. Lagrimas, M.E. Renz, L.G. Presta, D. Jackson, K.J. Hillan, S. Lew, and S. Fong. 1996. Mucosal addressin cell adhesion molecule-1: a structural and functional analysis demarcates the integrin binding motif. J Immunol. 157:2488–2497.

Wu, M., G. Chen, and Y.P. Li. 2016. TGF-beta and BMP signaling in osteoblast, skeletal development, and bone formation, homeostasis and disease. Bone Res. 4:16009.

Yan, D., M. Wiesmann, M. Rohan, V. Chan, A.B. Jefferson, L. Guo, D. Sakamoto, R.H. Caothien, J.H. Fuller, C. Reinhard, P.D. Garcia, F.M. Randazzo, J. Escobedo, W.J. Fantl, and L.T. Williams. 2001. Elevated expression of axin2 and hnkd mRNA provides evidence that Wnt/beta –catenin signaling is activated in human colon tumors. Proc Natl Acad Sci U S A. 98:14973–14978.

Yan, Y., D. Tang, M. Chen, J. Huang, R. Xie, J.H. Jonason, X. Tan, W. Hou, D. Reynolds, W. Hsu, S.E. Harris, J.E. Puzas, H. Awad, R.J. O’Keefe, B.F. Boyce, and D. Chen. 2009. Axin2 controls bone remodeling through the beta-catenin-BMP signaling pathway in adult mice. J Cell Sci. 122:3566–3578.

Yost, C., M. Torres, J.R. Miller, E. Huang, D. Kimelman, and R.T. Moon. 1996. The axis-inducing activity, stability, and subcellular distribution of beta-catenin is regulated in Xenopus embryos by glycogen synthase kinase 3. Genes Dev. 10:1443–1454.

Yue, R., B. Shen, and S.J. Morrison. 2016. Clec11a/osteolectin is an osteogenic growth factor that promotes the maintenance of the adult skeleton. Elife. 5.

Zhou, B.O., R. Yue, M.M. Murphy, J.G. Peyer, and S.J. Morrison. 2014. Leptin-receptor-expressing mesenchymal stromal cells represent the main source of bone formed by adult bone marrow. Cell Stem Cell. 15:154–168.

